# A global multiregional proteomic map of the human cerebral cortex

**DOI:** 10.1101/2021.03.07.434254

**Authors:** Zhengguang Guo, Chen Shao, Yang Zhang, Wenying Qiu, Wenting Li, Weimin Zhu, Qian Yang, Yin Huang, Lili Pan, Yuepan Dong, Haidan Sun, Xiaoping Xiao, Wei Sun, Chao Ma, Liwei Zhang

## Abstract

The Brodmann area (BA)-based map is one of the most widely used cortical maps for studies of human brain functions and in clinical practice; however, the molecular architecture of BAs remain unknown. The present study provided a global multiregional proteomic map of the human cerebral cortex by analyzing 29 BAs. These 29 BAs were grouped into 6 clusters based on similarities in proteomic patterns: the motor and sensory cluster, vision cluster, auditory cluster and Broca’s area, Wernicke’s area cluster, cingulate cortex cluster, and heterogeneous function cluster. We identified 474 cluster-specific and 134 BA-specific signature proteins whose functions are closely associated with specialized functions and disease vulnerability of the corresponding cluster or BA. The findings of the present study could provide explanations for the functional connections between the anterior cingulate cortex and sensorimotor cortex and for anxiety-related function in the sensorimotor cortex. The brain transcriptomic and proteomic comparison indicated that they both could reflect the function of cerebral cortex, but showed different characteristics. These proteomic data are publicly available at the Human Brain Proteome Atlas (www.brain-omics.com). Our results may enhance our understanding of the molecular basis of brain functions and provide an important resource to support human brain research.

## Introduction

The human cerebral cortex, the outer layer of the cerebrum, exhibits great complexity in its histological structure and cellular organization ^1^. In 1909, Korbininan Brodmann distinguished a total of 52 subareas based on the differences in cytoarchitectural organization across the cortex; these areas were later termed Brodmann areas (BAs) ^2^. Subsequently, BA-based map has become one of the most widely used cortical maps for the investigation of the brain functions and in clinical practice. BAs are often correlated with certain cortical functions. For example, Broca’s speech and language area is consistently localized in BAs 44 and 45. cytoarchitecture ^1, 2^, imaging ^1, 3^, and functional roles ^3^ of BAs have been extensively studied, and comprehensive human brain MRI imaging ^1^, histology ^1^, and connectome maps ^3^ have been investigated.

Molecular annotations, especially at the protein level, of various BAs will provide a deeper understanding of the molecular basis for diverse brain functions. Transcriptomic and proteomic maps of the brain have been generated to study the molecular basis of distinct cytoarchitecture and functions of various BAs ^1^. The Allen Institute for Brain Science has generated a transcriptional atlas of the human brain by comprehensive profiling of nearly 900 anatomically precise subdivisions using microarrays ^1^. The human neocortex is characterized by a relatively homogeneous transcriptional pattern; the molecular topography of the neocortex is similar to its cytoarchitecture. At the protein level, several studies generated a protein atlas of the mouse brain and identified some brain region-restricted proteins ^4, 5^ or cell type-specific proteins ^4^. In 2014, Kim *et al* drafted a proteomic profile of the human frontal cortex from a pool of three individuals and identified 9,060 proteins ^6^. In 2017, Becky el al. ^7^ performed a proteomic survey of 7 postnatal human brain regions, including two cerebral cortex regions, the cerebellar cortex, the hippocampus, and other neurological nuclei, ranging from early infancy to adulthood. Individual variations due to age and sex were considerably lower than variations between brain regions. Importantly, the differences in brain cytoarchitecture, development, and function are better represented by changes at the protein level than by changes at the transcriptional level.

Previous omics studies have increased our understanding of the brain function. However, some important scientific aspects have not been investigated. Functional annotation of the brain demonstrated that several brain regions are functionally connected but are not spatially adjacent, such as BA41/42 (auditory cortex) and BA 44/45 (Broca’s area and language production) ^3, 8^. However, the data on the brain transcriptome showed a relatively homogeneous expression pattern in the neocortex, and molecular topography in this area is similar to spatial topography. Proteins are the main functional operators in all cells. Proteomic map of the cerebral cortex may provide new insight into this scientific context. Previous studies of the brain proteome investigated only a few BAs, which are not sufficient to illustrate these aspects. Thus, a comprehensive proteomic map of various BAs of the human cerebral cortex is urgently needed.

The present study used 29 BA postmortem specimens to generate a global multiregional proteomic map of the human cerebral cortex. Initially, an extensive proteomic library of BA46 was generated. Then, 29 BAs were quantitatively analyzed using the 2DLC/MS/MS approach combined with isobaric tags for relative and absolute quantitation (iTRAQ) labeling. Clustering analysis of 29 BAs was performed based on the similarities in protein expression patterns. Cluster- and BA-signature proteins were identified and analyzed using bioinformatics. The major functional brain areas, including the cingulate cortex, motor cortex, sensory cortex, language area, and visual cortex, were functionally annotated using proteomic data. The transcriptomes of above BAs were also analyzed and the comparison of proteome and transcriptome on BA functions was presented. The results of the present study provide a molecular basis for BA functions. Furthermore, the database provides an important resource to support brain research.

## Results

### A broad and comprehensive atlas of brain proteome

A single human brain specimen was obtained from the Human Brain Bank facilitated by the China Human Brain Bank Consortium. The brain specimen was collected from a 54-year-old willed male donor who died of lung cancer and did not have any history of brain metastasis, trauma, or other diseases. The samples of 29 BAs of the left cerebral hemisphere were collected with a postmortem delay of less than 6 hours. Hematoxylin-eosin staining (HE staining) of the samples showed that the brain was cancer-free (Supplementary Figure 1). The protocols for brain tissue acquisition, dissection, and sample preparation were under consistent and stringent quality control (see Materials and Methods) based on the standardized operational protocol (SOP) of the Human Brain Bank of China ^9^.

BA46 (dorsolateral prefrontal cortex (DFC)) played an important role in many cognitive processes and had been comprehensively characterized in previous proteomic studies ^6, 7^. Therefore, deep proteomic profiling of BA 46 was performed using the 2DLC/MS/MS approach (Figure 1A) to construct a Chinese reference brain protein library. A total of 8,600 proteins (Supplementary Table 1 A-B, data quality in Supplementary Figure 2A) were identified in the present study, corresponding to 73% and 89% of the proteins identified in the studies of Kim *et al* ^6^ and Becky *et al*. ^7^, respectively. Additional 979 (11.4%) proteins were identified by brain proteome analysis for the first time (Figure 1B and Supplementary Table 1 A-B). Thus, we generated a broad and high-coverage proteomic dataset with identification depth similar to that achieved by the two most comprehensive datasets reported in previous studies^6, 7^.

**Figure 1:**
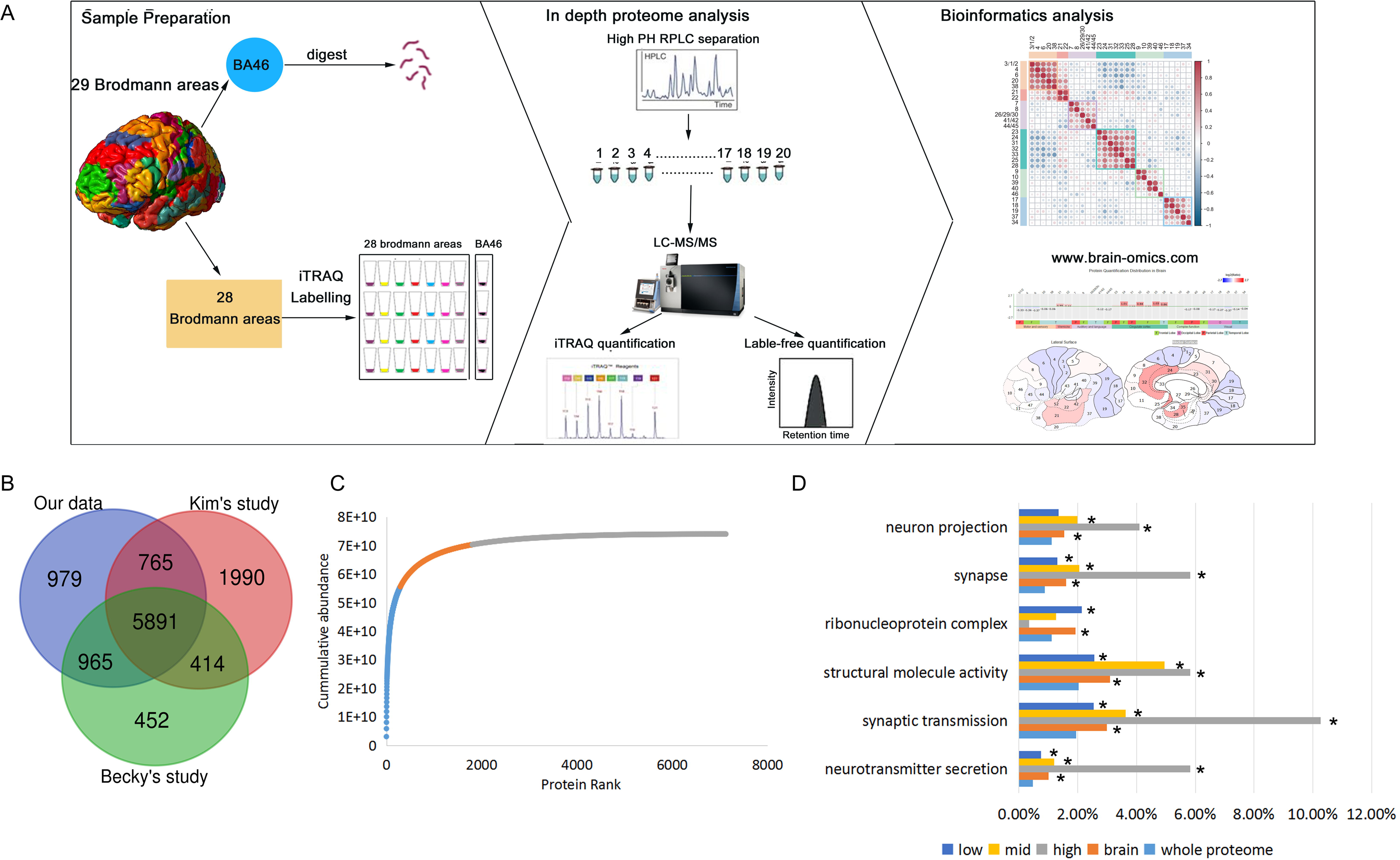
A broad and comprehensive atlas of brain proteome covering 29 BAs. **A:** Graphical illustration of the workflow for brain proteome database in Brodmann Area (BA46) and brain proteomic map in the 29 major BAs. First, an extensive proteomic library was generated of BA46by 2D-LC-MS/MS and label-free quantification in BA46. Then, 29 BAs were quantitatively analyzed by 2D-LC-MS/MS and iTRAQ quantification. Further, a website, Human Brain Proteome Atlas, HBPA(www.brain-omics.com) was built. B: Venn diagram of protein number of BA46 in present and previous studies. C: Accumulation of protein mass from the highest abundance to the lowest abundance protein in BA46. The high abundance proteins, (grey, top for 75% of total protein abundance), mid abundance proteins, (orange, top 75%-95% of total protein abundance), and low abundance proteins (blue, bottom 5% total protein amount) shown by distinct colors. D: GO enrichment analysis of the brain proteome, and the high, middle, low abundance brain proteins, and compared with the whole proteome. The percentages of the proteins in each category were in bar plot. *: p<0.05.

Then, to provide a comprehensive protein abundance analysis, the proteome abundance in BA46 was calculated by the intensity-based absolute quantification (iBAQ) method ^10^. A total of 7,129 proteins were successfully quantified, and their abundances spanned 7 orders of magnitude (Supplementary Figure 3). Approximately 4% of the most abundant proteins (292 proteins) accounted for 75% of the total protein abundance and were considered high-abundance brain proteins in the present study. The remaining 20% and 5% of the total protein abundances corresponded to 1,513 mid-abundance proteins and 5,324 low-abundance proteins, respectively (Figure 1C, Supplementary Table 1C). Gene ontology (GO) analysis revealed that nervous system structure and function were enriched in this brain proteome, especially in the case of high-abundance proteins (Figure 1D, Supplementary Table 1D), implying the important role of high-abundance proteins in brain functions.

Finally, to investigate specific proteomic profiling in BAs and explore the associations between BA protein expression and function, we used iTRAQ-based quantitative proteomics to generate a region-resolved proteome map of the brain covering 29 BAs with important significance for research and clinical applications (Figure 1A and Table 1). A total of 4,308 proteins were quantified in all BAs. The batch effect of 4 batches was evaluated by qualitative and quantitative analysis. The median CV of four batches was around 6% (Supplementary Figure 2A), and the Venn diagram of protein identification in four batches (Supplementary Figure 2B) showed high overlapping. The log ratio distributions of 29 iTRAQ quantification results(Supplementary Figure 2C) showed similar normal distribution, and the detailed ratio distributions of each BA in four batches (Supplementary Figure 2D) showed similar distributions.

**Table 1:**
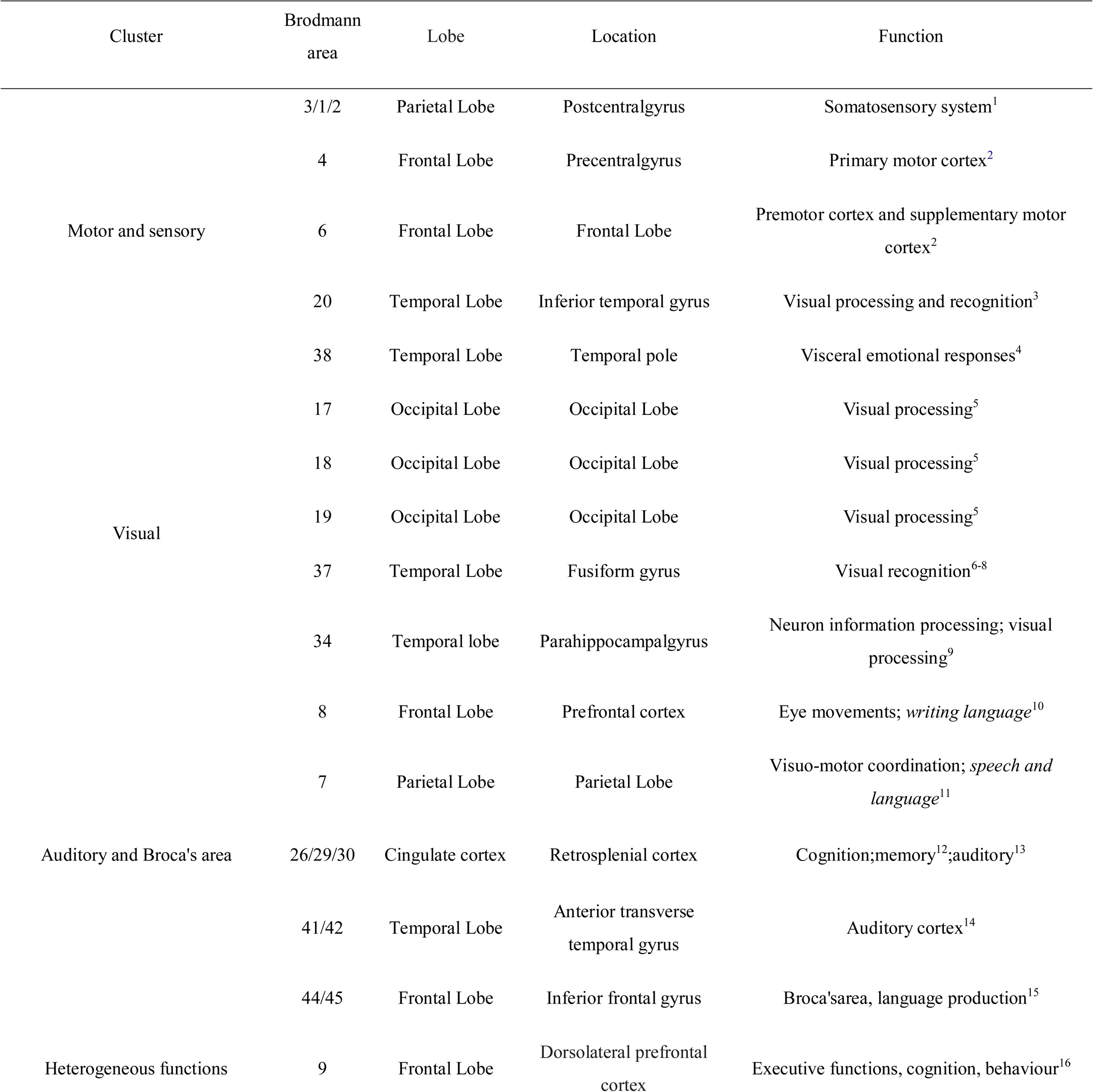
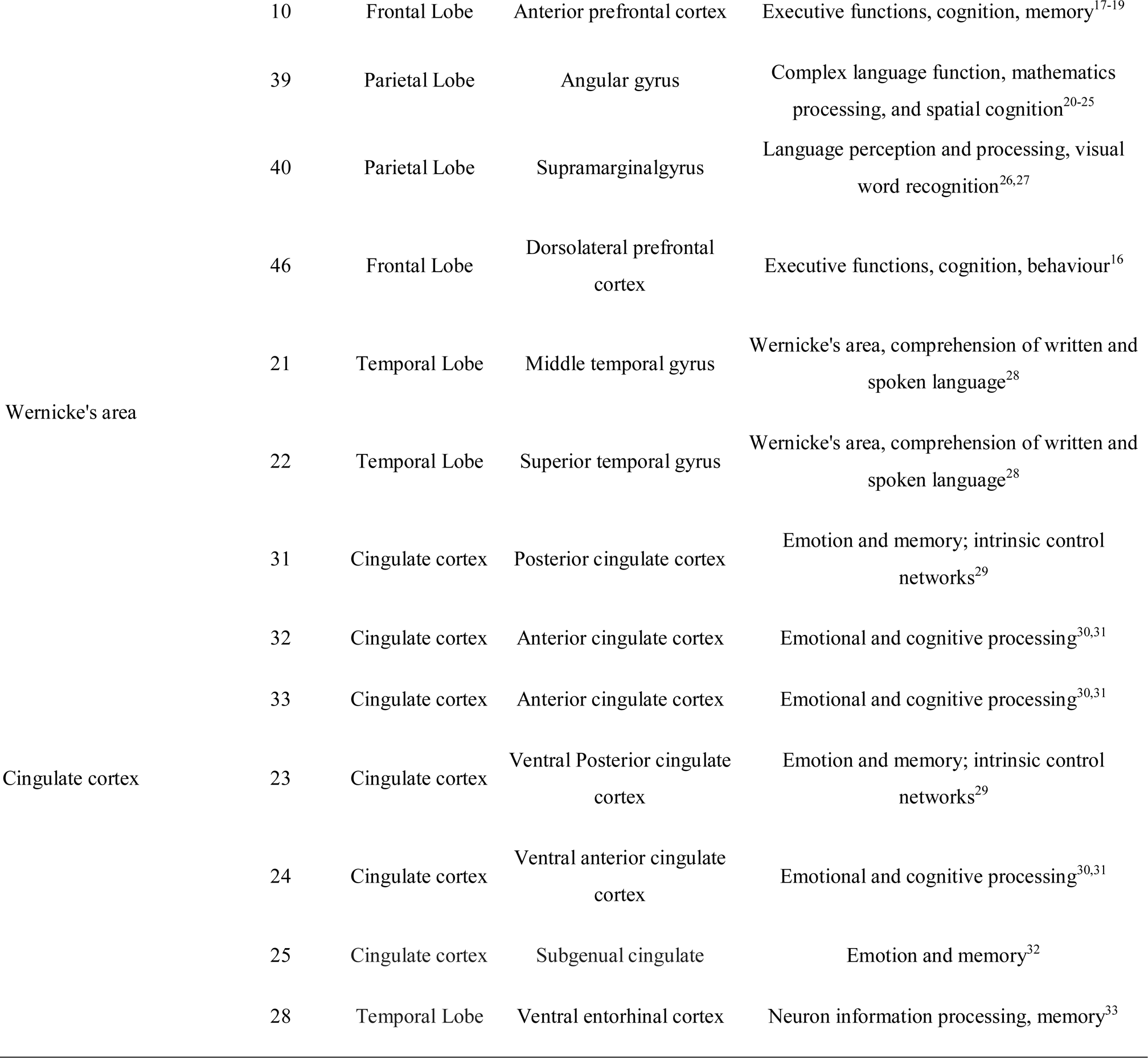
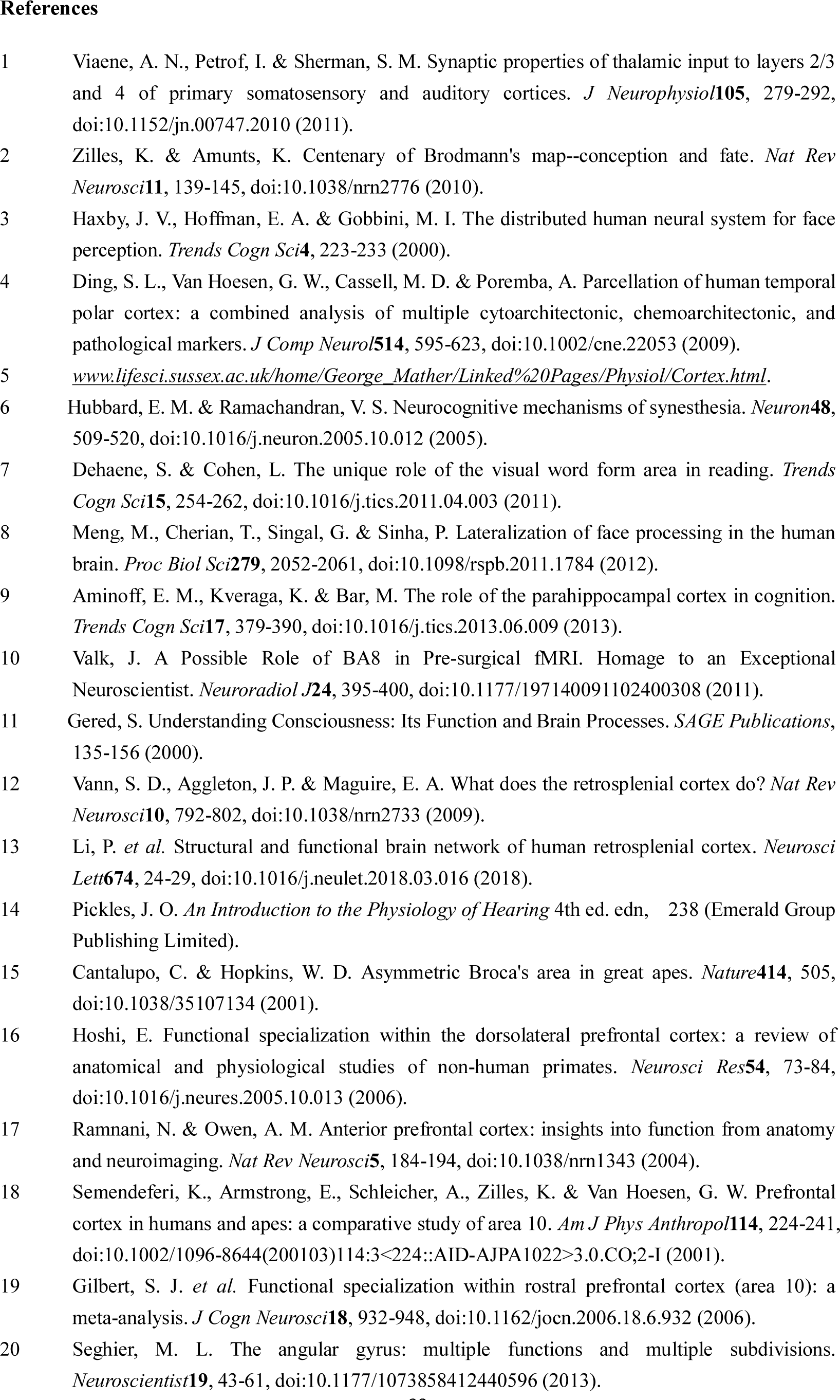
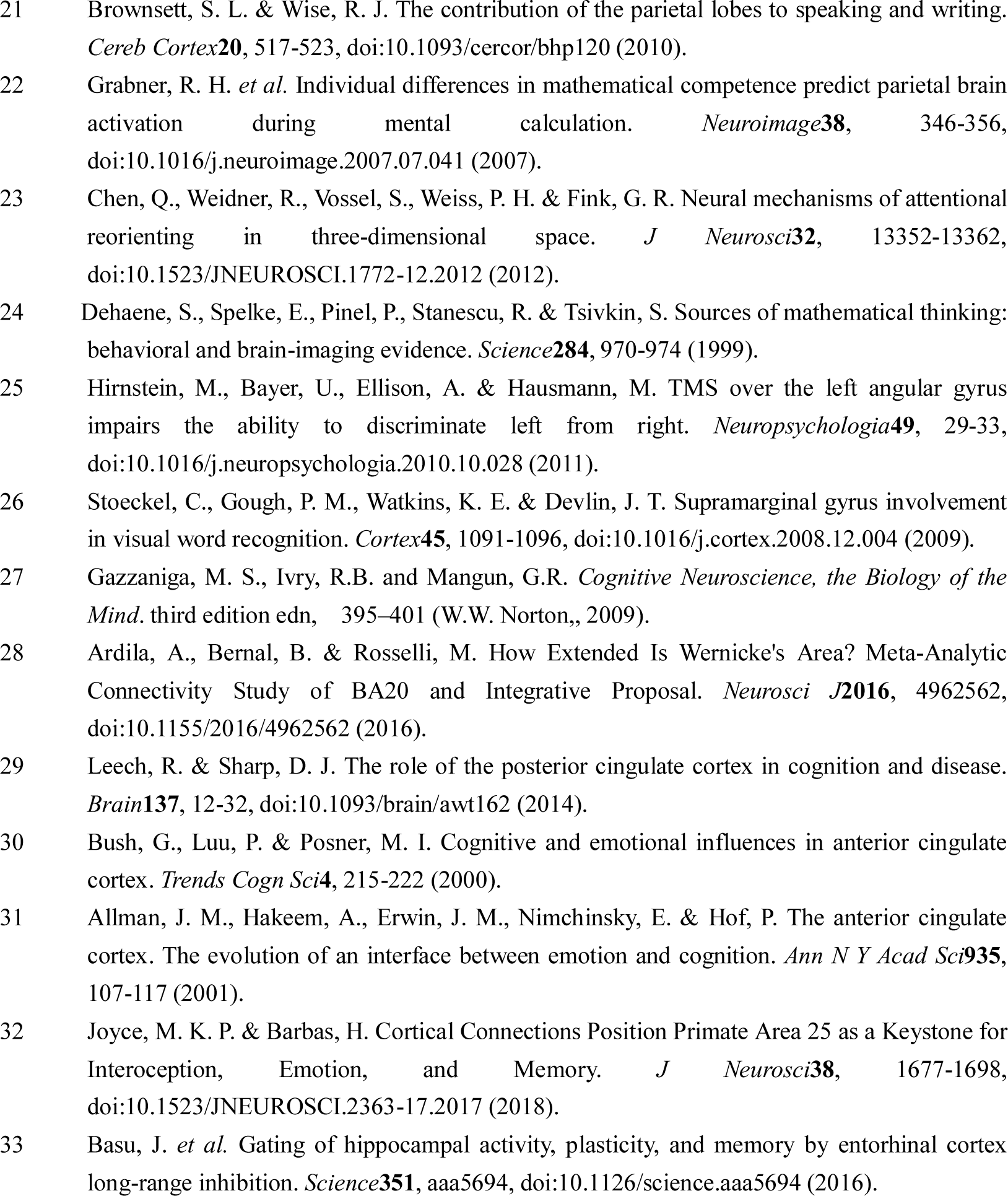
Location, function and proteomic clustering of 29 human brain Brodmann areas.

A total of 1,241 proteins were characterized by interarea variability (the difference between the maximum and minimum ratios ≥ 2, detailed in Method section). The Human Brain Proteome Atlas(www.brain-omics.com) was constructed to enhance visualization and use of these data, which are publicly accessible (Supplementary Figure 4).

### Brain proteomic atlas reflects functional parcellations

Proteins are the main components and functional operators in the cells; thus, Bas with similar protein expression patterns may have similar cytoarchitecture or perform similar functions. Based on the similarities of protein expression patternsK-means clustering^11^was used for the classification of BAs. The consensus cumulative distribution function (CDF) plot and silhouette plot (**Supplementary Figure 5**) based on the consensus matrix were used to group 29 BAs into 6 clusters (detailed data in Methods section, Figure 2A, and Table 1).

**Figure 2:**
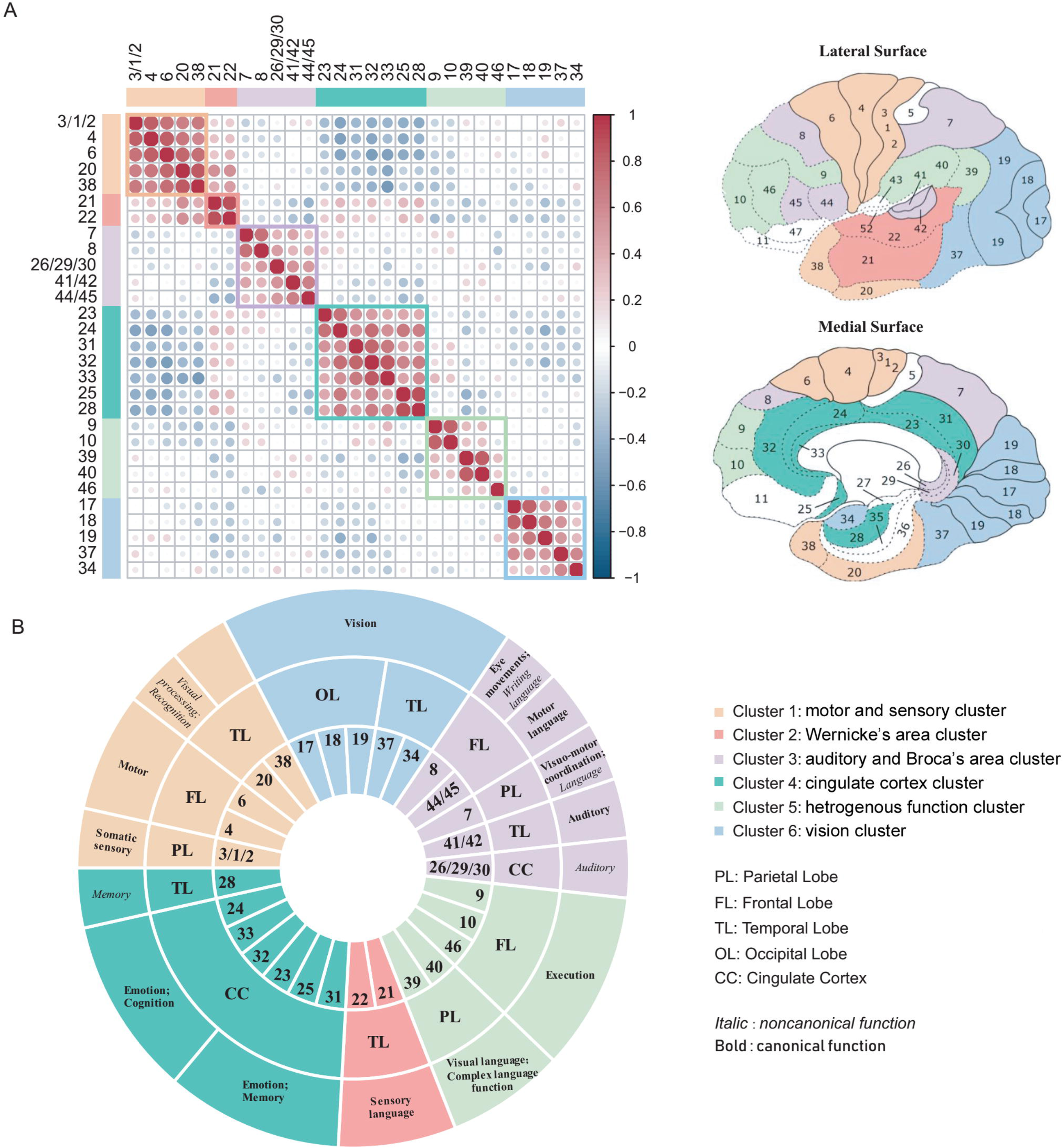
Brain proteomic atlas reflects the functional parcellation. A: The 29 brain areas were divided into 6 clusters, including motor and sensory cluster, vision cluster, auditory and Broca’s area cluster, Wernicke’s area cluster, cingulate cortex cluster, and heterogeneous function cluster. The pearson correlation heatmap of 29 BAs in the left. Red: positive correlation; Blue: negative correlation. The locations of 29 BAs were shown in the right. B: Brief annotations of 29 BAs in the 6 clusters. The lobe location of each BA was annotated in the inner circle, and the function of each BA was annotated in the outer circle. Bold: Canonical function; Italic: uncanonical function.

The cingulate cortex cluster (cluster 4) included BAs in the cingulate cortex, subgenual cingulate cortex, and entorhinal cortex. All these BAs belong to the transition areas between the allocortex (3 layers of neuronal cell bodies) and neocortex (6 layers of neuronal cell bodies). Therefore, the cytoarchitecture of the cingulate cortex cluster was different from that of the neocortex ^12^. At the proteome level, the cingulate cortex cluster was characterized by protein patterns distinct from those of other clusters, which were mainly located in the neocortex (Figure 2A), implying that the brain proteome can reflect the differences in cytoarchitecture between the cingulate cortex and the neocortex.

The remaining 23 BAs within the neocortex were grouped into 5 clusters, and BAs within a single cluster were functionally similar. For example, the vision cluster (cluster 6) comprised a group of BAs (BAs 17, 18, 19, 37, and 34) engaged in visual processing (Figure 2B, Table 1). BAs 17, 18, and 19 are located in the occipital lobe, and BA37 is adjacent to BA19. All four BAs correspond to classic visual processing regions^13–15^. BA34 is located in the parahippocampal gyrus and is not spatially connected to other four BAs, although BA34 was reported to be related to visual processing^16^. Another example is the auditory and Broca’s area cluster, which is composed of several spatially separated but functionally related BAs (Figure 2B). BA44/45 of this cluster (Broca’s area and motor language area), BA8 (writing language area) ^17^ and BA7 (engaged in speech ^18^) are directly involved in language processing, whereas BA41/42 (auditory cortex) and BA26/29/30 (engaged in sound processing) ^19^ are associated with auditory processing, which is indirectly linked to language processing ^3, 8^.

Interestingly, two language processing BAs, BA22 (classic Wernicke’s area and sensory language area) and adjacent BA21 (also considered a part of Wernicke’s area ^20^), were clustered into the Wernicke’s area cluster. This cluster was characterized by proteomic patterns distinct from the auditory and Broca’s area cluster. To illustrate the differences in protein expression between these two clusters, we compared differential proteins between the two clusters. A total of 88 and 108 proteins were expressed at higher levels in the Wernicke’s area cluster and auditory and Broca’s area cluster, with a 1.5-fold change, respectively (Supplementary Table 3). Differentially expressed proteins were enriched in metabolic pathways, suggesting metabolic differences between the two language regions (Supplementary Figure 6A). These proteins are also related to neurotransmitter pathways ^21^, including glutamate, -aminobutyric acid (GABA), and dopamine pathways (Supplementary Figure 6B) ^21^. In particular, the GABA receptor was expressed at a higher level in the auditory and Broca’s area cluster compared to that in the Wernicke’s area cluster in agreement with previous reports on an important role of the GABA receptor in the physiological functions of Broca’s area. An inhibitor of the AGABA receptor causes a dysfunction in Broca’s area and induce stuttering ^22^, and a GABA agonist improves chronic Broca’s aphasia^21^. Therefore, these differentially expressed proteins provide valuable information about the intrinsic mechanism of language processing.

### Cluster/BA signature proteins linked to specialized function of clusters/BAs

BAs with similar protein expression patterns tend to have similar cytoarchitecture and functions; thus, cluster signature proteins (proteins that were expressed at a high level in a certain cluster) may provide information about specialized functions of a cluster. Iterative signature analysis (ISA, details described in the Methods section)^23^ was used to identify a total of 474 cluster signature proteins in the present study (Figure 3A and Supplementary Table 4A). Pathway, function, and disease annotations of these proteins are shown in Supplementary Table 4B-E, Figure 3B-3C, and Supplementary Figure 7A-7B.

**Figure 3:**
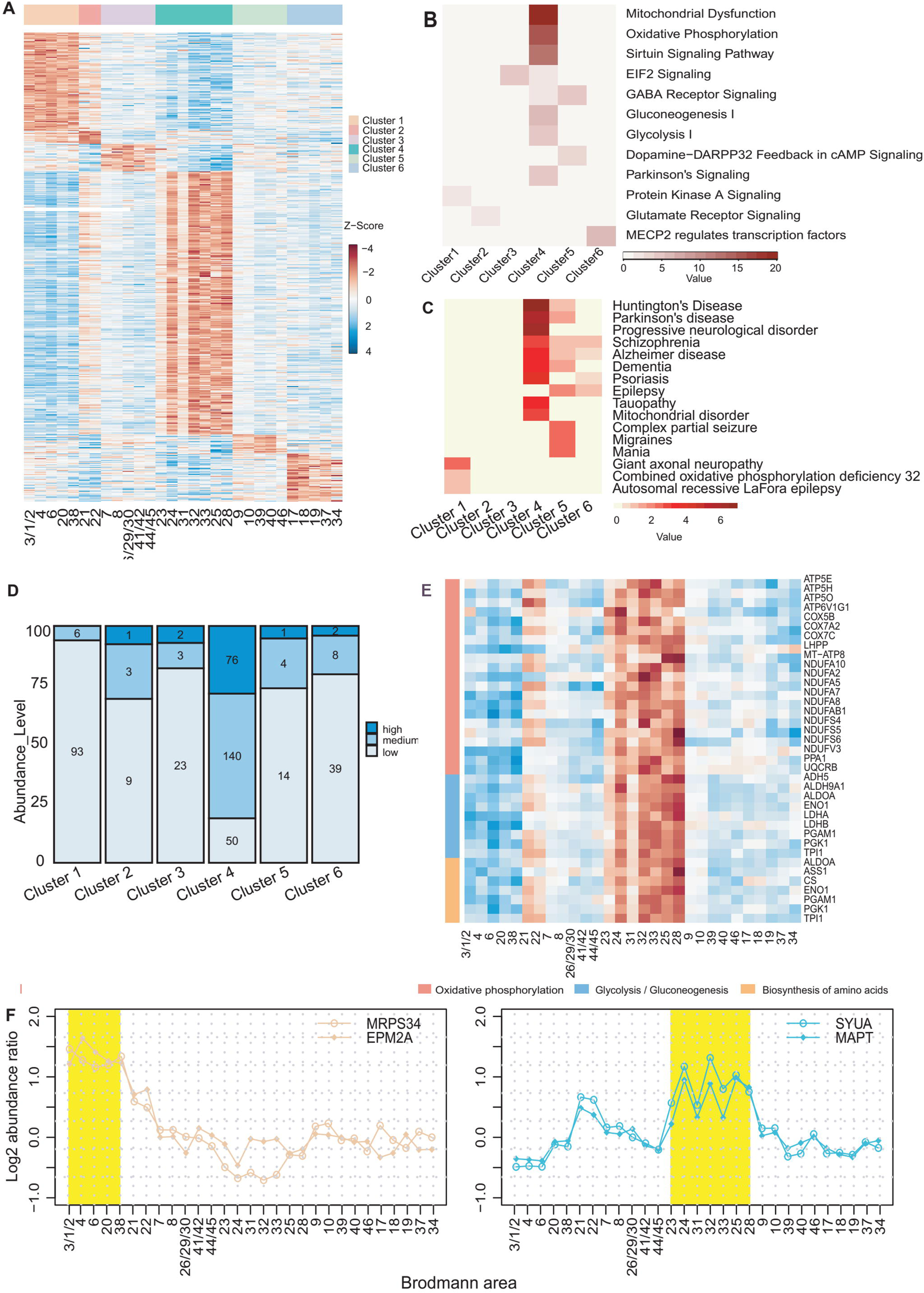
Cluster signature proteins linked to specialized function of the cluster. A: Heat map of the cluster signature proteins. The normalized protein expression value z-score represents the protein abundance. Red: high abundance. Blue: low abundance. B,C: Pathway (B) and disease (C) enrichment analysis of cluster signature proteins in six clusters. The −Log (p-value) by two-sided hypergeometric test were in heatmaps.Red:significantly enriched. D: Distribution of the high-, mid-, low-abundance proteins in the cluster signature proteins. The number of high-, mid-, low-abundance proteins in each cluster signature proteins were annotated in the bar plot. E: Heatmap of cingulate cortex cluster signature proteins involved in glycolysis/gluconeogenesis, oxidative phosphorylation, and biosynthesis of amino acids. The normalized protein expression value z-score represents the protein abundance. Red: high abundance. Blue: low abundance. F: The relative expression value of two motor and sensory cluster signature proteins (MRPS34 and EMP2A) and two cingulate cortex cluster specific proteins (SYUA and MAPT) in all the 29 BAs. The motor and sensory cluster and the cingulate cortex cluster were highlighted by yellow in the left and right panels, respectively.

The cingulate cortex cluster had the largest number of signature proteins, 266 out of all 474 signature proteins. These proteins included 76 (26.3%) and 140 (52.6%) high- and mid-abundance proteins, respectively (Figure 3D and Supplementary Figure 7C-7E), corroborating the results regarding the differences in cytoarchitecture and protein expression between the cingulate cortex cluster and the neocortex. Functional analysis showed that these signature proteins are enriched in metabolic pathways, including important enzymes for oxidative phosphorylation (NADH dehydrogenase), glycolysis (ALDOA, PGK1, PGAM1, and ENOA), glycogen turnover (TPIS), and amino acid synthesis (ASS1 and PDHX) (Figure 3E), indicating high metabolic activity of the cingulate cortex. These results are in agreement with the metabolic analysis of the cingulate cortex reported in a previous study ^24^. A total of 21 signature proteins were involved in cognition, emotion, and memory (Supplementary Table 4F), consistent with the functional characteristics of the cingulate cortex ^24–26^. These results are in agreement with the functional characteristics of the cingulate cortex. Disease annotations indicated that proteins related to neurological degenerative diseases (Huntington’s disease, Parkinson’s disease, and Alzheimer’s disease) were enriched in the signature proteins of the cingulate cortex cluster. Several key proteins associated with signaling in Alzheimer’s and Parkinson’s diseases were expressed at high levels in the cingulate cortex cluster (Supplementary Figure 7F-7G). These proteins include tau (MAPT) and alpha-synuclein (SYUA), which are important biomarkers for Alzheimer’s disease ^27, 28^ and Parkinson’s disease ^29^, respectively (Figure 3F). The pathological accumulation of these two proteins is etiologically correlated with the development of Alzheimer’s ^28^ and Parkinson’s diseases ^29^. The data of Western blot analysis showed that the protein expression patterns of MAPT and SYUA were consistent with the patterns detected by proteomic analysis (Supplementary Figure 8A-B). The results of IHC-Fr also demonstrated that MAPT and SYUA were expressed at a high level in BA25 compared to that in BA4 (Supplementary Figure 8C). The results of this study demonstrated the regional distribution of MAPT and SYUA for the first time, adding new information about pathogenesis and vulnerability of these two neurodegenerative diseases in the cingulate cortex. Furthermore, another brain (brain 2) was used to validate the cluster-specific signature proteins by parallel reaction monitoring (PRM). Detailed clinical information on brain 2 is provided in the Methods section. PRM analysis validated that 28 signature proteins were expressed at a high level in cingulate cortex cluster (Supplementary Table 5A-5B).

The motor and sensory cluster had 99 signature proteins. These proteins were enriched in the protein kinase A (PKA) pathway (Figure 3B). The PKA pathway is associated with brain motor function (neuronal plasticity in motor neurons ^30, 31^) and sensation (pain sensitization in sensory neurons ^32, 33^). Striatal-enriched protein tyrosine phosphatase (STEP), a molecule downstream of PKA, is associated with motor skill learning in the motor cortex^34^. Other signature proteins with potentially interesting functions include the neuron survival protein EMP2Aandmitochondrial 28S ribosomal protein S34(MRPS34) (Figure 3F). The functional loss of EMP2A was reported to cause sensorimotor cortex excitability in Lafora disease ^35^. MRPS34 mutation causes delayed psychomotor development and neurodevelopmental deterioration in Leigh syndrome^36^. Immunological analysis confirmed that EMP2A and MRPS34 were expressed at a high level in the motor and sensory cluster (Supplementary Figure 8). Two signature proteins of motor and sensory cluster, EMP2A and PTN5, were validated by PRM (Supplementary Table 5A-5B).

The vision cluster included 49 signature proteins enriched in the methyl-CpG-binding protein 2 (MECP2)-regulated transcription factor pathway. MECP2 and transcriptionally regulated transcription factors are involved in neurological system and neuropsychiatric disorders ^37^ (Figure 3B). MECP2 plays an important role in normal development of the central nervous system and maintenance of neurological functions. Mutation of the MECP2 gene leads to Rett syndrome, which is an autism spectrum-associated disorder with visual impairment^38^. In a mouse model, knockout of MECP2 causes progressive dysfunction of the visual cortex and rapid regression of visual acuity ^39^. Three signature proteins of vision cluster, MECP2, TAGLN, and CAVN1, were confirmed to be expressed at high level by PRM (Supplementary Fig 10B, Supplementary Table 5A-5B). Additionally, Becky et al ^7^ reported a proteomic comparison of BA17 (primary visual cortex) and BA46 (DFC) in 11 individuals ranging from early infancy to adulthood. By comparing the 49 vision cluster signature proteins in present study with Becky’s study, 16 were expressed at high level in primary visual cortex in both studies (Supplementary Table 5E).

Additionally, we focused on BA signature proteins (protein abundance in one or two BAs that is more than twofold higher than the median abundance) to investigate BA-specific functions. A total of 134 BA signature proteins were identified based on the analog criteria suggested by Uhlen*et al* ^40^ (details are provided in the Methods section), (Figure 4A and Supplementary Table 4G). These BA signature proteins included several proteins closely associated with the corresponding brain functions. In the cingulate cortex, superoxide dismutase [Cu-Zn] (SOD1) was identified as a signature protein in the anterior cingulate cortex (BA32 and BA24). SOD1 mutation interferes with GABAergic neurotransmission, leading to cortical neuronal loss in this brain area^41^. In the neocortex, FAM107B was identified as a signature protein in BA41/42 (auditory cortex) (Figure 4B). Knockout of FAM107B leads to hearing impairment in mice^42^. Three BA signature proteins, transgelin-2 (BA17 and BA18), RAD23A (BA40), and FAM107B (BA41/42), were validated by Western blot (Figure 4B, Supplementary Figure 9). Five BA signature proteins, NEFH (BA4), PMP2 (BA4), CALB2 (BA28), TAGLN (BA17 and BA18), and FAM107B (BA41/42), were validated to be expressed at a high level in corresponding BAs in brain 2 (Supplementary Table 5C-5D). Two primary visual cortex signature proteins, TAGLN and H1F0 were also found to be expressed at high level in primary visual cortex in Becky’s study^7^(Supplementary Table 5E).

**Figure 4:**
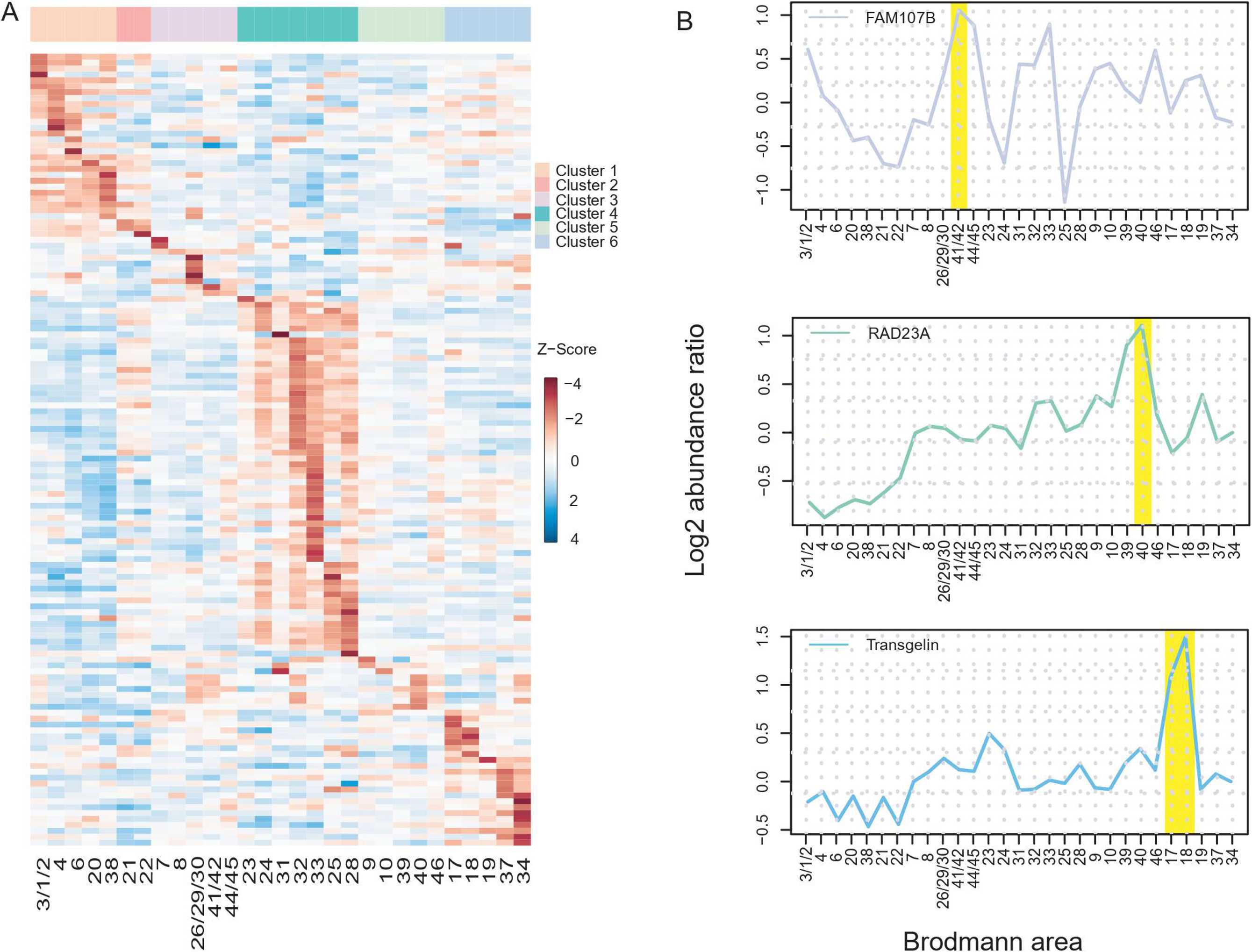
BA signature proteins linked to BAs’ function. A: Heat map of the BA signature proteins in 29 BAs. The normalized protein expression value z-score represents the protein abundance. Red: high abundance. Blue: low abundance. B: The relative expression value of three BA signature proteins, TAGLN, RAD23A, and FAM107B in all the 29 BAs. The BA41/42, BA40, BA17 and BA18 were highlighted by yellow in the upper, middle, and lower panels, respectively.

Overall, these findings demonstrated intrinsic links between specialized functions of the brain clusters and BAs and molecular functions of the corresponding signature proteins, which may help to understand the mechanisms of physiological or pathophysiological functions of the brain.

### Molecular annotation of uncanonical functions of BAs

#### Functional connection between the motor cortex and anterior cingulate cortex

The anterior cingulate cortex is tightly connected to the motor cortex through neuronal cell projections according to the results of fluorescence labeling imaging in rats^43^. In humans, Caruana*et al*. demonstrated that the anterior cingulate cortex plays important roles in the control of complex motor behaviors according to the results of intracorticalelectrical stimulation ^44^. Molecular evidence of the connection between these two functional brain regions requires additional investigation.

Proteomic data obtained in the present study demonstrated that a series of proteins were expressed at a high level in the primary motor cortex (BA4) and anterior cingulate cortex (BA33) (Figure 5A, Supplementary Table 6A). Most of these proteins are associated with motor neuron structure and function (Figure 5B). Neurofilament proteins (NEFL, NEFM, NEFH, and INA) are the main components of motor neurons^45^ and are necessary for the maintenance of the number, morphology, viability, and regenerative capability of motor neurons ^46–48^. Myelin basic protein promotes myelination of motor neurons ^49^. MIF ^50^ and PVALB ^51^ are associated with amyotrophic lateral sclerosis, a neurodegenerative disease targeting motor neurons ^52^. Thus, these proteins may represent molecular basis of the functional connections between the primary motor cortex (BA4) and anterior cingulate cortex (BA33) and may also play a role in the motor functions of the anterior cingulate cortex.

**Figure 5:**
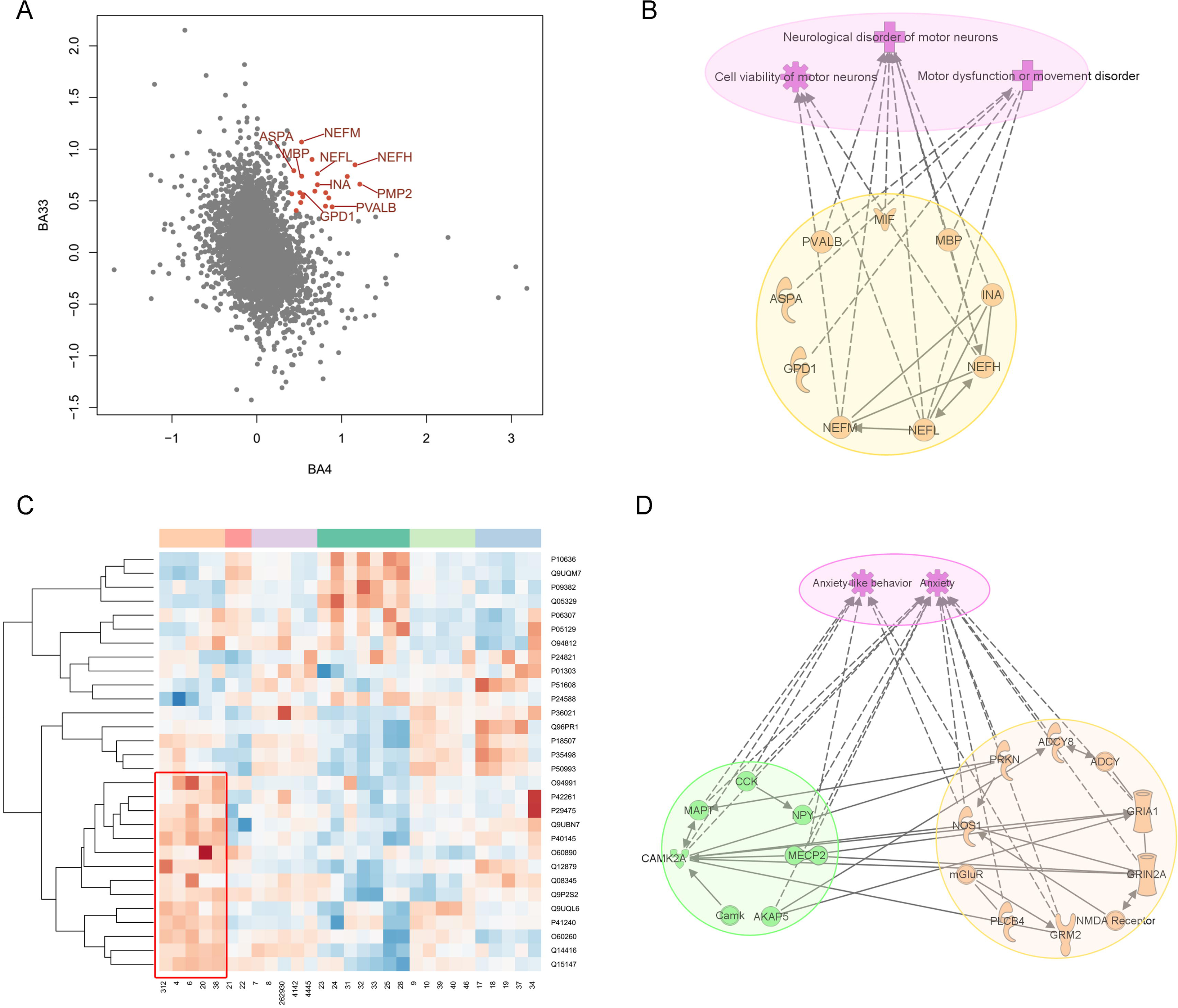
Molecular annotation of the un-canonical functions of BAs. A: Scatter plot of protein log2 fold expression value in BA 33 compared with the value in BA4. Proteins expressed at high level in both BA4 and BA33 highlighted in orange. B: The co-highly-expressed proteins in BA4 and BA33 were associated with motor neuron structure and functions. C: Clustering analysis of anxiety-related proteins. The proteins highly expressed in motor and sensory cluster highlighted in red. D: Protein network of anxiety-related proteins. Orange: The proteins expressed at high level in motor and sensory cluster. Green:The proteins expressed at low level in motor and sensory cluster.

#### The motor and sensory cluster may be involved in anxiety

Anxiety disorder is a disease characterized by excessive and persistent worry and fear and is very common worldwide (lifetime prevalence is 5∼25% of the population), representing a substantial economic burden ^53, 54^. The sensorimotor cortex has been reported to be associated with social anxiety disorder ^55^ and anxiety-related personality traits ^56^ according to the results of electrophysiological studies. However, the molecular basis of anxiety in the sensorimotor cortex requires further investigation.

Proteomic data obtained in the present study indicated that 30 of the 1,241 proteins with inter area variability are involved in anxiety or anxiety-like behaviors according to IPA (Supplementary Table 6B). Clustering analysis indicated that nearly half (14, 46.6%) of these 30 proteins were expressed at a high level in the motor and sensory cluster (Figure 5C). Some of these 14 proteins, including ADCY8, GRIA1, GRIN2A, PLcB4, SLITRK5, NOS1, and GRM2, formed a protein interaction and regulation network and participated in the expression of anxiety and anxiety-like behaviors (Figure 5D). PlcB4 ^57^ and NOS1 ^58^ were expressed at a high level in the medial septum and the hippocampus (two anxiety behavior-related regions), respectively, and inhibition of these genes in the regions with high levels of expression increased anxiolytic effects in a mouse model. GRIN2A and GRIA1 are two important glutamate receptors involved in the function of the sensorimotor cortex ^59^ and also involved in anxiety. Knockout of GRIN2A ^60^ and GRIA1 ^61^ causes a decrease in anxiety in a mouse model. Thus, we hypothesized that these proteins are expressed at a high level in the motor and sensory cluster and may be involved in anxiety-related functions in the sensorimotor cortex, which may explain the molecular basis of anxiety-related processes in the sensorimotor cortex.

#### Transcriptomic characterization of brain BAs

The transcriptomes of brain BAs were explored to reveal the characteristics of the brain functions. 29 BAs by RNA-seq were analyzed, and 21 samples passed the quality control (RIN>=6, 28S/18S>=0.7, gene mapping>=40%) and were used for further functional analysis. An average of 17,137 transcripts was detected based on the RNA-seq data (Supplementary Table 7A, Supplementary Figure 10). Comparison of the brain proteome library and the transcriptomic database of BA46 identified 8,411 transcriptomic entries (97.8%) for 8,600 proteins detected by LC-MS (Figure 6A). The proteomic data covered 48.8% of the transcriptomic data, covering 61% with expression level>1 read per kilobase per million mapped (RPKM) and 78.4% with expression level>10 RPKM (Figure 6B).

**Figure 6:**
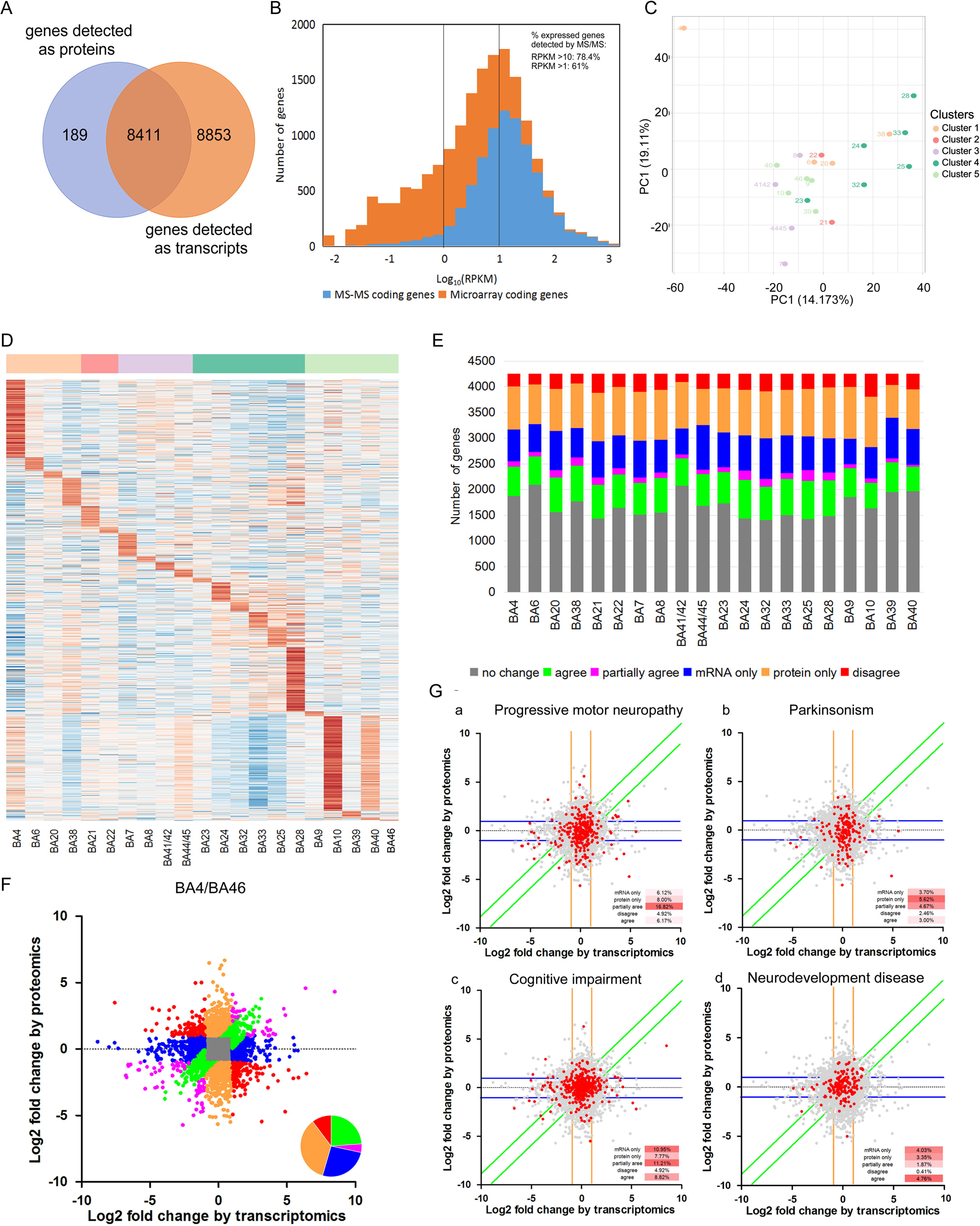
Comparison of brain proteome and transcriptome. A: Venn diagram of protein and mRNA number in BA46. B: Histogram of the distribution of the gene expression in mRNA and protein level. Orange: Gene identified in protein level. Grey: Genes only identified only in mRNA level. The MS/MS coding genes covered 61% genes expressed at>1 read per kilobase per million mapped (RPKM) and 78.4% of genes expressed at >10 RPKM. C: Principal component analysis of mRNA shows that the BA4 and cluster cingulate cortex were partially separated with the other BAs. D: Heat map of the BA signature genes in 27 BAs. The normalized protein expression value z-score represents the protein abundance. Red: high abundance. Blue: low abundance. E: The Distribution of the relative dominance of no change (grey), agree (green), partially agree (purple), protein only (Blue), mRNA only (orange), and disagree (red) genes. Genes are defined based on the consistency or inconsistency between the mRNA and protein measurements. Agree genes are those with same direction of change in both RNA and protein level, and with the same magnitude of fold change (the variability in protein level between BAs is < 2 fold of that in mRNA level). Partially agree genes are genes with same direction of change, but with different magnitude of fold change (larger than 2 fold) in RNA and protein level. No change genes, both protein and RNA were not changed (within 2 fold) between the BAs. Protein only genes vary between different BAs by protein (larger than 2 fold), but not by mRNA (within 2 fold). mRNA only genes vary by mRNA (larger than 2 fold), but not by protein (within 2 fold). Disagree genes performed opposite direction of change between protein and mRNA. F:Scatter plots of Log2 formed mRNA and protein abundance fold change between BA4 and BA46. grey (no change), Green (agree), purple (partially agree), Blue (protein only), orange (mRNA only), and Red (disagree) genes were defined in E. The inset pie charts illustrated the percentage of the green, purple, blue, orange, and red genes. G: Scatter plots of mRNA and protein fold change for BA4/ BA46 comparison, showing the position of gene products contained in ontological terms of interest. The inset small table showed the percentage of the genes of responding ontological terms in the mRNA only, protein only, agree, partially agree and disagree categories. The colored lines indicated the threshold of mRNA gene (orange), protein only gene (blue), and the agree gene (green).

A total of 17,091 genes were quantified in 21 BAs. PCA of the transcriptomic data showed that BA4 (primary motor cortex) was different from other BAs in agreement with the finding of the Allen Institute that a series of genes are specifically enriched in the primary motor cortex^1^. The cingulate cortex cluster tended to be partially separated from other BAs, which was partially consistent with the proteomic results. However, four functionally distinct clusters distinguished by proteomic analysis could not be found based on the transcriptomic data (Figure 6C).

The BA signature genes at the transcriptomic level were used to explore BA-specific functions. A total of 1,319 BA signature genes were identified (Figure 6D and Supplementary Table 7B). Functional analysis indicated that BA signature genes were closely involved in the functions of corresponding BA. In cingulate cortex, a total of 13 signature genes of BA 28 were involved in memory function, in agreement with the functional characteristics of the ventral entorhinal cortex (BA28) ^26^(Supplementary Table 7C). In neocortex, motor neuron related proteins (NEFH, NEFM) ^46,47^were identified as signature genes of primary motor cortex (BA4), and were also expressed at high level in BA4 in our proteomic data. In Auditory cortex(BA41/42), SMPX was identified as a signature gene. Previous study reported that mutation of SMPX was associated with X-linked deafness type 4 ^62^.

A total of 4,250 genes were quantified based on the data of transcriptomic and proteomic results in 21 BAs (Supplementary Table 7D). Gene products for all region pairs were classified based on fold-change similarity between mRNA and protein (details described in the Methods section). An average of 39.5% of the genes were classified into “no change” (≤ 2-fold for both protein and RNA) category, and 17.4% of the genes were classified into “agree” category (same direction of changes (>2-fold) and same magnitude of fold changes) or “partially agree” category (same direction of changes (> 2-fold) and different magnitude of fold changes) category. In addition, 20.6% or 15.7% of the genes were considered “protein only” (≥2-fold change in protein level only) or “RNA only” (≥2-fold change in mRNA level only), respectively. In contrast, 6.7% of the genes corresponded to opposite differences in the mRNA and protein levels (> 2-fold) (“disagree” group) (Figure 6E).

To show the differences between transcriptome and proteome on brain functions, BA4 (Primary motor cortex) and BA46 (DFC) were compared and the scatter plots of the mRNA and protein fold change between them were shown in Figure 6F (Supplementary Table 7E, other BA pair comparison data in Supplementary Figure 11). According to IPA functional annotation of the differential gene products between BA4 and BA46, Progressive motor neuropathy and Parkinsonism related genes were enriched in “protein only” and “partially agree” categories, which could reflect the characteristics of the Primary motor cortex. In contrast, the cognitive impairment and the depressive disorder related genes showed a bias toward“mRNA only” categories, reflecting the characteristics of the DFC Additionally, the neurodevelopment related genes were enriched in the “agree” category, consistent with the Becky’s findings ^7^ (Figure 6G, Supplementary Table 7F).

## Discussion

In this study, we constructed a Chinese cerebral cortex proteomic atlas of 29 BAs from a donor without neurological diseases. Similarities in the protein expression patterns indicated that these 29 BAs can be grouped into six clusters. The cluster- and BA signature proteins can provide molecular evidence for annotation of cytoarchitecture, function, and disease vulnerability of these clusters or the corresponding BAs. All data are publicly available in the Human Brain Proteome Atlas (HBPA) (www.brain-omics.com).

In this brain proteome atlas, BAs with similar protein expression patterns have similar functions, even though they are not spatially adjacent, which reflects functional parcellations. BAs within the cingulate cortex cluster and vision cluster are characterized by both spatial proximity and functional relevance. However, BAs within the auditory and Broca’s area cluster are not directly adjacent and are functionally connected (speech and auditory). These results indicated that the proteomic map mainly reflects the functional parcellations of the human cerebral cortex. Furthermore, proteomic similarity of BAs may provide molecular annotations for functional connections between BAs. For example, BA38 (temporal pole) is an enigmatic region with largely unknown function ^63^. The brain connectome indicated a functional connection between BA3/1/2-BA38 and BA21/22-BA38 ^3^. The results of fMRI experiments indicated that BA38 is involved in speech comprehension and considered a part of “extended Wernicke’s area” ^64, 65^. A study on Braille reading in blind individuals using brain imaging methods indicated thatBA38 can integratesensory-motor and language comprehension. ^63^ In the present study, BA38 was characterized by protein expression patterns similar to both the sensorimotor cortex (BA3/1/2, BA4, and BA6) in the motor and sensory cluster and Wernicke’s area (BA21 and BA22). We suggest that BA38 may play integratory roles in both sensory-motor functions and language comprehension.

Additionally, the present study identified 474 cluster- and 134 BA signature proteins. These proteins reveal intrinsic links between their molecular functions and specialized functions of the brain regions. The cingulate cortex contains an extremely large number of signature proteins compared to that of other neocortex regions. These signature proteins were enriched in metabolic pathways, mainly glycolysis, glycogen turnover, and amino acid synthesis. Previous studies demonstrated that astrocytes had unique metabolic pathways of brain glucose metabolism compared to those in neurons ^66^. First, glycolysis is predominant in astrocyte glucose metabolism. Second, the storage and turnover of glycogen are mainly concentrated in astrocytes. Third, TCA cycle-derived amino acids, such as aspartate (Asp), glutamate (Glu), glutamine (Gln), arginine (Arg), and γ-aminobutyric acid (GABA), are synthesized in astrocytes using astrocyte-specific enzymes ^66^. The combination of previous reports and proteomic data obtained in the present study indicates the activation of astrocyte metabolism in the cingulate cortex, which may correspond to metabolic characteristics of the cingulate cortex. Additionally, the results of the present study suggest that some signature proteins in the cingulate cortex cluster participate in the pathogenesis of some neurological diseases, such as MAPT in Alzheimer’s disease and SYUA in Parkinson’s disease. Previous studies have shown that the accumulation of these two proteins in the cingulate cortex is a pathological feature of the corresponding diseases ^28, 29^. Abnormalities in the cingulate cortex play a central role in the development of Alzheimer’s ^24^ and Parkinson’s diseases ^67, 68^. These signature proteins may be investigated to determine molecular mechanisms of neurological diseases, especially in some specific brain regions. Thus, information about these proteins may substantially improve our understanding of the molecular basis of brain functions and pathogenesis.

Furthermore, the data of the present study provide molecular annotations of the noncanonical functions of brain regions. Molecular function analysis of these data provides molecular basis for functional connections between the anterior cingulate cortex and motor cortex and may explain the molecular basis of anxiety in the sensorimotor cortex. These findings promote our understanding of the molecular mechanisms of the noncanonical functions of brain regions. Therefore, proteomic data obtained in the present study may be used to investigate new functions of BAs or new functional connections between BAs.

In present study, the comparison of BA transcriptome and proteome indicated thatthe proteomic data could separate the cerebral cortex into 6 clusters, which were associated with BA’s function. However, transcriptomic data did not show similar classification, which was consistent with Allen results^1^. Above results indicated that brain proteomic data might preferably reflect BA function. Functional analysis of the trancriptome suggested that it could also reflect the function characteristics of BAs. 17.4% gene showed similar differential expression trend as the protein (such as neurodevelopment), and some BA signature genes (NEFH, NEFL in primary motor cortex) also showed consistent expression patterns as the protein. Moreover, 15.7% gene only differentially expressed in mRNA level, and some of which were associated with BA function (cognition impairment, depressive disorder in DFC). Above data demonstrated that transcriptome and proteome showed different characteristics of brain functions. According to previous studies ^7^, in neuron mRNA and proteins might locate in different brain regions. Synapse proteins were transmitted from the nucleus into another BA through axon, so they could be detected in both BAs. However, mRNA was limited to the source BA containing cell body, so they were only identified in source BA. Thus, such genes would be differentially expressed only on mRNA level. Above study could partially account for the differences between transcriptome and proteome.

This study established a Chinese proteome atlas of the brain; however, the following issues need to be resolved in the future studies. First, we drafted a global multiregional proteomic map of the left hemisphere of a single brain. Previous proteomic and transcriptomic studies suggested that individual variations in age and gender were substantially lower than variations across brain regions ^1, 7^. Proteomic atlas generated in the present study can represent common characteristics of the normal human cerebral cortex to some extent; however, future studies should consider inclusion of multiple individuals balanced with regard to gender, age, and brain hemisphere sampling and should use more sensitive and accurate proteomic technologies. Second, this pilot atlas includes only 29 BAs with important brain functions, and future studies should include additional important regions, such as the cerebellar cortex, hippocampus, and other important nerve nuclei, to generate a more comprehensive atlas.

## Materials and methods

### Tissue procurement

Samples from frozen postmortem human brain specimens were obtained following the SOP of the Human Brain Bank of China facilitated by the China Human Brain Bank Consortium (see Qiu*et al*. ^9^ for details on sample collection, handling, and preservation). Basic demographic information, medical information, and medical records of the donor were obtained. Brain 1 specimens were collected from a 54-year-old willed male donor who died of lung cancer and did not have any history of brain metastasis, trauma, or other diseases. Brain 2 was collected from a 93-year-old willed female donor who died of heart failure and did not have neurological diseases. The scalp was incised along the coronal plane from the ear to top of the skull and turned over to expose the skull. The skull was cut apart at the sagittal suture starting 1 cm above the eyebrows and continuing to the occipital protuberance, and the dura was removed. The cerebrum, brain stem, cerebellum, and part of the cervical cord were completely removed and temporarily preserved in cold saline. Twenty-nine samples of the cortex from various BAs were dissected from the left cerebral hemisphere and stored at −80°C for protein extraction. Routine hematoxylin/eosin, silver, and immunohistochemical staining procedures for amyloid-beta, p-tau, and alpha-synuclein were performed in the visual cortex, inferior parietal lobule, superior and middle temporal gyri, medial frontal cortex, amygdala, anterior cingulate gyrus, hippocampus, basal ganglia, cerebellum, middle brain, cerebral pons, and medulla in the right cerebral hemisphere of the brain sample; the data were used for pathological diagnosis to exclude neurodegenerative diseases. According to the Standardized Operational Protocol ^9^, we performed HE staining of the brain areas, including BA40, 24, 17, 41/42/22, to confirm that brain is cancer-free.

### Ethical statements

Brain tissue was collected after obtaining consent from the donor and the next of kin using a signed informed consent form and the project was approved bythe Institutional Review Board of the Institute of Basic Medical Sciences, Chinese Academy of Medical Sciences (approval number: 009-2014).

### Sample preparation

A total of 29 brain tissue samples were selected for proteomic analysis (80 mg each). The samples were rinsed with PBS until washing fluid was clear, and each sample was lysed with lysis buffer (containing 8 M urea,2.5 M thiourea,4% CHAPS, 65 mM DTT, and 50mMTris-HCl) in a homogenizer on ice. The lysate was centrifuged at 14,000 × rpm at 4°C for 20 min, and the supernatant was collected. The Bradford method was used to measure the protein concentration. Proteins were digested by the FASP method, as previously described^69^. The peptides were extracted by an Oasis C18 extraction column and quantified by the BCA method.

### iTRAQ labeling and 1D off-line separation

Digested peptides were labeled with 8-plex iTRAQ reagent according to the manufacturer’s protocol (Sciex, Framingham, MA). The 28 Brodmann areas were randomly divided into 4 groups (Supplementary Table2) and labeled byiTRAQ8-plex agent. In each group, BA46 was used as common internal reference sample. The labeled samples within each experiment were mixed together. To reduce sample complexity, the peptide samples were fractionated through a high-pH RPLC system. The samples were reconstituted in 100 µL of buffer A(1 mM ammonia, pH= 10) and loaded onto a column (4.6 mm×250 mm, Xbridge C18, 3 μm, Waters, Milford, MA). The elution gradient was composed of 5–25% buffer B (90% ACN, 1 mM ammonia, pH = 10) ataflow rate of0.8 mL/min for 60 min.60 fractions were collected, dried, andresuspended in 0.1% formic acid; the fraction were pooled into 20 samples using a stepwise concatenation strategy to combine every 20^th^ fraction (1, 21, 41; 2, 22, 42;…). A total of 100 fractions from a group of unlabeled samples and four groups of iTRAQ-labeled samples and were further analyzed by LC-MS/MS.

### LC-MS/MS analysis

The pooled mixtures of **iTRAQ-**labeled samples were analyzed using a self-packed RP C18 capillary column (75 μm×100 mm, 1.9 μm). The elution gradient was composed of 5–30% buffer B (0.1% formic acid and 99.9% ACN) at a flow rate of 0.3 μL/min for 45 min. An LTQ Orbitrap Fusion Lumos instrument (Thermo Fisher Scientific, Waltham, MA) was used to acquire raw data. The following parameters were used for MS data acquisition: top speed data-dependent mode (3 s) was used for full scan; full scans were acquired in an Orbitrap at a resolution of 60,000; the MS/MS scans were acquiredat a resolution of 15,000 with charge state screening (excluding precursors with unknown charge state or +1 charge state); 38% normalized collision energy in HCD mode; 30s dynamic exclusion (exclusion size list 500);1.6 Da isolation window. Each fraction from the iTRAQ-labeled sample was run twice. For label-free analysis, 32% normalized collision energy in HCD was used, other parameters were identical to the iTRAQ. Each fraction from the unlabeled sample was run three times.

### Database search

The MS/MS spectra were searched by Proteome Discoverer software (v3.1, Thermo Fisher Scientific) using the SwissProt human database downloaded from the UniProt website (www.UniProt.org). Trypsin was selected as the cleavage specificity parameter, andallowed a maximum of two missed cleavagesites. **For iTRAQ quantification**, carbamidomethylation of cysteine and iTRAQ 8-plex labels were set as fixed modifications, and the oxidation of methionine, deamidation of asparagine and glutamine, and carbamylation of lysine and peptide N-terminus were used as dynamic modifications. The searches were performed using a peptide tolerance of 10 ppm and a production tolerance of 0.02 Da. A 1% false-positive rate at the protein level was set as a filter, and each protein had to contain at least 1 unique peptide. **For label-free quantification**, carbamidomethylation of cysteine was used as a fixed modification, and other search parameters were identical to the parameters used for iTRAQ quantification. The abundance of each protein was estimated by Progenesis software (Nonlinear Dynamics, version 4.0), and the iBAQ value of each protein was calculated based on the abundance of a protein divided by the theoretical polypeptide number of the corresponding protein.

### Bioinformatics analysis of the iTRAQ data of 29 BAs

Each iTRAQ experiment contained seven BA samples and a common internal reference sample (BA46). In each experiment, the abundance ratios of the seven samples in BA46 were calculated, and the average protein abundance ratios of two technical replicates were used for further analysis. To ensure reliability of the data, the missing values in any two technical replicates of any group were excluded from subsequent analysis. The abundance ratios of proteins identified in all four experiments were merged into a single table in which each row represented a protein and each column represented a BA. In the column for BA46, all protein ratios were filled with 1. Then, the ratios were log2 transformed and normalized to a column median of 0 and a row median of 0. Thus, the positive value was higher than median, and the negative value were lower than median. Through row-wise normalization, the expression pattern of each row/protein in all the samples could be better exhibited, and it would be helpful for following cluster analysis and signature proteins filtering.

For better cluster analysis and signature protein selection the proteins without inter-area variability were filtered. The fold changes of all proteins between all BA pairs were calculated. The distribution of log2 fold change showed that the and Q3 + 1.5 interquantile range was 1.57. A strict filter (fold change > 2) was used as the threshold of outliers, which would be helpful to signature proteins. To identify the differential proteins between Wernicke’s area cluster and auditory and Broca’s area cluster, a relative relax threshold (1.5 fold change) was used to find more differential proteins, which would be helpful for pathway or function enrichment analysis.

Consensus clustering was performed using the Consensus Cluster Plus package of R^11^ to assess the similarities in the proteome profiles of BAs. K-means clustering with Euclidean distance was performed 1,000 times on the subsampled data. The optimal number of clusters was selected using the consensus matrix, consensus cumulative distribution function (CDF) plot, and silhouette plot. The consensus matrix showed the best clustering results at k = 3. However, the CDF of consensus increased at greater k values. At k = 6, a relative increase in CDF reached a very small value. The silhouette plot at k = 6 showed a mean silhouette score of 0.72, and all scores were positive, indicating that clusters were well separated. Thus, BAs were clustered into six clusters.

Cluster signature proteins were identified by a biclustering algorithm known as iterative signature algorithm (ISA). ISA was implemented using the isa2 package for R^23^. Biclustering was restricted to upregulated modules in both rows and columns with a score threshold of 0.7. For all six clusters, modules that included at least 80% BAs inside a cluster and no BAs outside of the cluster were selected. Proteins in the selected modules with scores that passed the threshold of 0.7 were considered signature proteins for this cluster.

An adequate cutoff ratio was determined to identify proteins overexpressed in specific BAs. The differences in the log2 protein abundance ratios between all BA pairs for all proteins were calculated, and the data were normally distributed with a mean of 0.00 and a standard deviation (sd) of 0.35. The log2-fold change cutoff was then determined to be 1, which approximated the mean + 3 * sd. Thus, an area signature protein was defined as a protein with abundance level in one or two particular BAs of at least twofold of the median abundance level in all 29 BAs.

### Downstream Bioinformatics analysis

All proteins of interest were analyzed using the David database (http://david.ncifcrf.gov/) and compared with the whole human genome for GO enrichment analysis. For IPA analysis, the signature or differential proteins were evaluated using IPA software (Ingenuity Systems, Mountain View, CA), and the enrichment of the disease, function, and canonical pathway categories were analyzed according to the P-value rankings. For reactome analysis, cluster signature proteins were mapped to the reactome database in the pathway category according to the P-value rankings. For KEGG analysis, cluster signature proteins were mapped to the KEGG database in the pathway category according to the P-value rankings.

### IHC and Western blot

Western blot analysis of brain samples was performed to validate the proteomic quantitation of several selected candidate proteins (tau, alpha-synuclein, EPM2A, MRPS34, TAGLN, FAM107B, and RAD23A). Proteins extracted from the brain tissues were separated by SDS-PAGE and electro-transferred to a PVDF membrane (Millipore, Bedford, MA). The membrane was blocked with 2% (v/v) BSA for 2 h at room temperature and incubated with primary antibodies and a peroxidase-conjugated secondary antibody. The membranes were visualized using Immobilon Western chemiluminescent horseradish peroxidase substrate (Millipore), and the bands were analyzed by ImageJ software.

Formalin-fixed, paraffin-embedded brain tissue samples were used for IHC-Fr analysis. The tissue sections were deparaffinized and rehydrated in xylene and a graded ethanol series, and antigen retrieval was performed in a pressure cooker by boiling in sodium citrate buffer (pH 6.0) for 2 min. Then, endogenous peroxidases were blocked with 0.3% H_2_O_2_ for 15 min, and the samples were incubated in the presence of 2% fetal calf serum for 20 min. Primary antibodies were incubated at 4°C overnight, and peroxidase-labeled polymer conjugated to anti-mouse, anti-rabbit, or anti-goat immunoglobulins was incubated for 1 h. The sections were counterstained with Mayer’s hematoxylin and dehydrated, and the images were acquired under a microscope.

Primary antibodies for Western blot and IHC-Fr against MAPT (rabbit monoclonal, ab76128), SYUA (rabbit monoclonal, ab138501), EPM2A (rabbit monoclonal, ab129110), MRPS34 (mouse monoclonal, HPA042112, Sigma), **TAGLN** (rabbit polyclonal, ab14106), and RAD23A (mouse monoclonal, ab55725) were purchased from Abcam (Cambridge, UK). Antibodies against FAM107B (rabbit polyclonal, ab122566) were purchased from Sigma-Aldrich (St Louis, MO).

### PRM analysis

Selected cluster and BA signature proteins were validated in 27 BAs of brain 2 by PRM. To estimate the quality of the data, we used the mixed sample as quality control (QC) during the analysis, before and after all samples, and every 7-8 samples. Two technical repeats of each sample were assayed. The average Pearson correlation coefficient of QC samples was 0.97, indicating the reproducibility of the QC.

The samples were analyzed on a C18 RP self-packed capillary LC column (75 μm×100 mm). The elution gradient was composed of 5-30% buffer B (0.1% formic acid, 99.9% ACN) with a flow rate of 0.5 μL/min for 45 min. An LTQ Orbitrap Fusion Lumos instrument was used to acquire raw data. Following parameters was used for MS data acquisition: PRM mode; full scans and MS/MS scans were acquired in Orbitrap at a resolution of 60,000 and 15,000, respectively; 32% normalized collision energy in HCD mode; 30s dynamic exclusion; The isolation window was 4, and the schedule mode window was 7 min.

PRM data processing was performed by Skyline 19.1 software as previous described ^69^. The peptide abundance was normalized to TIC in each sample, which was extracted by Progenesis QIP software (Waters).

### RNA seq and data processing

RNA-seq was performed by BGI cooperation (Shenzhen, China). For transcriptome data analysis, read counts were normalized using the median of ratios normalization method in DESeq2 to eliminate the variations in sequencing depth and RNA composition in the samples ^70^. Normalized counts were logarithmically transformed with base 2. A total of 17,091 transcripts with read counts > 1 in at least 50% of the samples were retained for downstream analysis. Transcriptomic and proteomic data were matched by gene symbols.

### Comparison between the proteome and transcriptome

The genes commonly quantified at the mRNA and protein levels were quantitatively compared. Both transcriptomic and proteomic datasets were log2 transformed by zero-mean normalization. The gene products for all possible region pairs were classified based on their fold-change similarity between mRNA and protein. Genes with the same direction of the changes at the RNA and protein levels and with the same magnitude of fold change (the variability in protein level between BAs was < 2-fold of that at the mRNA level, assigned to the “agree” category); the genes with the same direction of change but with different magnitude of fold change (larger than 2-fold) at the RNA and protein levels were assigned to the “partially agree” category. The genes that had no significant inter regional changes in either RNA or protein levels (within 2-fold) between BAs were assigned to the “no change” category. The genes that had variable levels of corresponding proteins between various BAs (larger than 2-fold) but did not vary at the mRNA level (within 2-fold) were assigned to the “protein only” category. The genes that had variable mRNA levels (larger than 2-fold) but did not vary at the protein level (within 2-fold) were defined as “mRNA” only. The genes that showed opposite changes in mRNA and protein levels were assigned to the “disagree” category.

## CRediT author statement

**Zhengguang Guo:** formal analysis, funding acquisition, validation, writing - original draft. Chen Shao: methodology, formal analysis, software visualization, funding acquisition, and writing - review & editing. Yang Zhang: conceptualization, writing - review &editing. Wenying Qiu: resources, investigation. Wenting Li: resources. Weimin Zhu: project administration. Qian Yang: validation. Yin Huang: software.Lili Pan: visualization.Yuepan Dong: visualization. Handan Sun: investigation. Xiaoping Xiao: investigation. Wei Sun: investigation, project administration, funding acquisition, writing - review & editing. Chao Ma: project administration, funding acquisition. Liwei Zhang: conceptualization, project administration, funding acquisition, supervision. All authors read and approved the final manuscript.

## Acknowledgments

We are grateful to Professor BaiLu, and Dr. He You from Tsinghua University for helpful discussions. This work was supported by National Key Research and Development Program of China (No. 2016 YFC 1306300, 2018YFC0910202), National Natural Science Foundation of China (No. 30970650, 31200614, 31400669, 81371515, 81170665, 81560121), Beijing Medical Research (No.2018-7), Beijing Natural Science Foundation (No. 7173264, 7172076), Beijing cooperative construction project (No. 110651103), Beijing Science Program for the Top Young (No. 2015000021223TD04), Beijing Normal University (No. 11100704), Peking Union Medical College Hospital (No. 2016-2.27), CAMS Innovation Fund for Medical Sciences (2017-I2M-1-009), the CAMS special basic research fund for central public research institutes (No. 2017PT310004), and Biologic Medicine Information Center of China, National Scientific Data Sharing Platform for Population and Health.

## Supplementary material

**Supplementary Figure 1: The HE staining data of two brains. A: Brain 1; B: Brain 2.**

**Supplementary Figure 2: Reliability of iTRAQ analysis pipeline.** A: Technical CV distributions of label-free and iTRAQ quantification. B: The venn diagram of the proteins quantified in 4 batches. C: Log ratio distribution of iTRAQ quantification results. D: The ratio distributions of each run in 4 batches.

**Supplementary Figure 3: Protein abundance distribution of BA46 from the highest abundance to the lowest abundance protein.**

**Supplementary Figure 4:Data visualization in Human Brain Proteome Atlas website.** A: Protein quantification distribution of MAPT in 29 Brodmann areas. B: Protein expression data in Brodmann area 25 (Subgenual area). C: Comparison of the differential proteins between BA3/1/2, and BA4.

**Supplementary Figure 5: The protein clustering analysis.** A: The clustering is derived by consensus clustering based on 1,000 resampled data sets, exploring the range of from k=2 to k=8. B: Based on both consensus cumulative distribution function (CDF) area curve and the delta plot assessing change in CDF area, the CDF of consensus increased at greater k values. At k = 6, a relative increase in CDF reached a very small value. C: Silhouette plots were performed to evaluate the Stability of the clustering. The silhouette plot at k = 6 showed a mean silhouette score of 0.72, and all scores were positive.

**Supplementary Figure 6: Analysis of the differential proteins between Wernicke’s area cluster and auditory and Broca’s area cluster.** A: IPA-based Canonical pathway analysis of differential proteins. The −Log (p-value) of the Canonical pathway term were shown. B: Heatmap of differential proteins involved in language eloquent regions related neurotransmitter pathways. The normalized protein expression value z-score represents the protein abundance. Red: high abundance. Blue: low abundance.

**Supplementary Figure 7: Functional analysis of cluster signature proteins, related to Figure 3.** A: The IPA-based biofunction (A) and KEGG database-based pathway (B) enrichment analysis of cluster signature proteins. The −Log (p-value) by two-sided hypergeometric test in heatmap. Red: significantly enriched. C-E: Heat map of the high-abundance proteins (C), mid-abundance proteins (D) and low abundance proteins(E) in the 29 BAs. The normalized protein expression value z-score represents the protein abundance. Red: high abundance. Blue: low abundance. F: Protein expression analysis of cingulate cortex cluster in Parkinson’s signaling. G: Protein expression analysis of cingulate cortex cluster in Alzheimer’s disease signaling. (Red: highly-expressed in cingulate cortex cluster; Green: lowly-expressed in cingulate cortex cluster)

**Supplementary Figure 8: The Western Blot and IHC-Fr results of cluster signature proteins.** A: Western Blot results of two motor and sensory cluster signature proteins (MRPS34 and EPM2A) and two cingulate cortex cluster signature proteins (Tau and a-syn). B: Comparison between immunoblotting results and iTRAQ results of above proteins. C: IHC-Fr validation results of two cingulate cortex cluster signature proteins (MAPT and SYUA). Scale bars represent as 50 µm.

**Supplementary Figure 9: The Western Blot and IHC-Frresults of Brodmann area signature proteins.** A:Western Blot results of three Brodmann area signature proteins (Transgelin, FAM107B, and RAD23A). B: Comparison between the immunoblotting results and iTRAQ results of above proteins.

**Supplementary Figure 10: Quality control of RNA-seq.** A: Number of gene detected in RNA-seq analysis of the 21 BAs. B: Box plot of gene expression level of 21 BAs.

**Supplementary Figure 11: Scatter plots of mRNA and protein fold change of all regions compared with BA 46**. These scatter plots are identically defined as those in Figure 6E.

**Supplementary Table 1: Brain proteome database in BA46.** A:Qualitative data of BA46 technical repeat 1, 2 and 3. The proteins identified in Kim’s study and Becky’s study were annotated. B: Protein identification summary of BA 46. C: Quantitative data of BA 46. D: The GO analysis results of brain proteome and high-, mid-, and low-abundance proteins compared with the whole human proteome.

**Supplementary Table 2: iTRAQ quantification data for 29 BAs.** A: Batch 1; B: Batch 2; C: Batch 3; D: Batch 4. E: Protein identification summary of the human brain proteome of 29 BAs. F: Normalized protein quantification data of the 4308 proteins in all of 29 BAs. The 1241 proteins with inter-area variability (the difference between the maximum and minimum ratios ≥ 2) were annotated.

**Supplementary Table 3: Differentially proteins between the Wernicke’s area cluster and Auditory and Broca’s area cluster.**

**Supplementary Table 4: Cluster/BA signature proteins.** A:List of cluster signature proteins. B: Biofunction analysis of cluster signature proteins. C: Disease analysis of cluster signature proteins. D: Canonical pathway analysis of cluster signature proteins. E: KEGG pathway analysis of cluster signature proteins. F: Cluster cingulate cortex signature proteins were involved in cognition, emotion, and memory. G: BA signature proteins.

**Supplementary Table 5: Validation of the Cluster/BA signature proteins. A:** PRM results of Cluster signature proteins (peptide level). B: PRM results of Cluster signature proteins (protein level). C: PRM results of BA signature proteins (peptide level). D: PRM results of BA signature proteins (protein level). E: Comparison of Vision cluster/vision cortex signature proteins results between this study and Becky’s study.

**Supplementary Table 6: Molecular annotation of the un-canonical functions of BAs.** A: Proteins highly expressed in both Primary motor cortex (BA4) and Anterior cingulate cortex (BA33). B:Thirty proteins with inter-area variability were involved in anxiety or anxiety-like behaviors.

**Supplementary Table 7. Comparison of brain proteome and transcriptome.** A:RNA-seq results for the 21 BAs. B: BA signature genes. C:Thirteen signature genes involved in memoryin BA28. D. The common genes in transcriptomic and proteomic results. E: The differential transcriptomic and proteomic results for BA4 and BA46 comparison. F. The disease and biofunction analysis of differential genes of BA4 and BA46. The categories mentioned in the manuscript were highlighted in yellow.

