## Supplementary Figure for "A global multiregional proteomic map of the human cerebral cortex"

**Supplementary Figure 1: The HE staining data of two brains. A: Brain 1; B: Brain 2.**

**A**

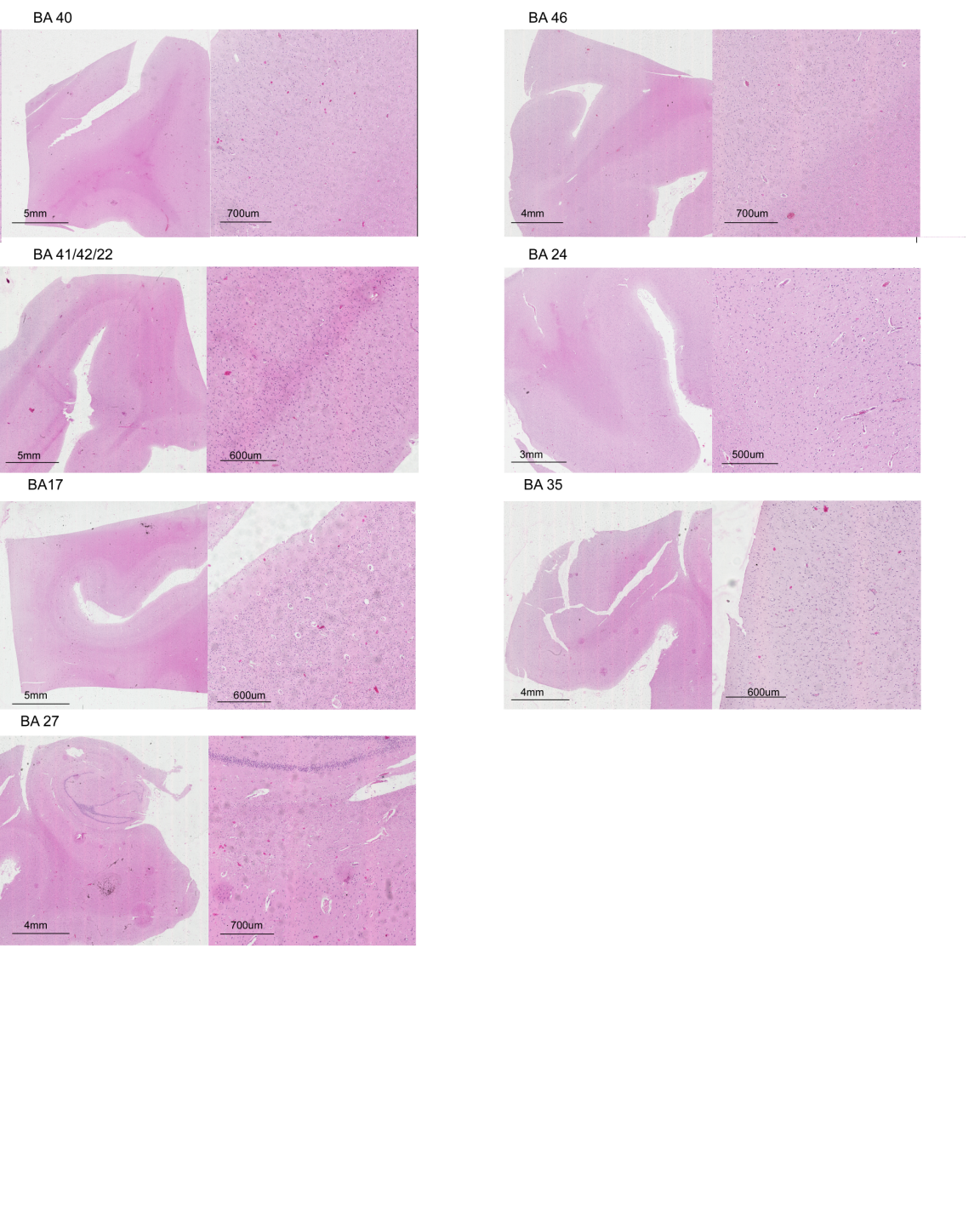

**B**

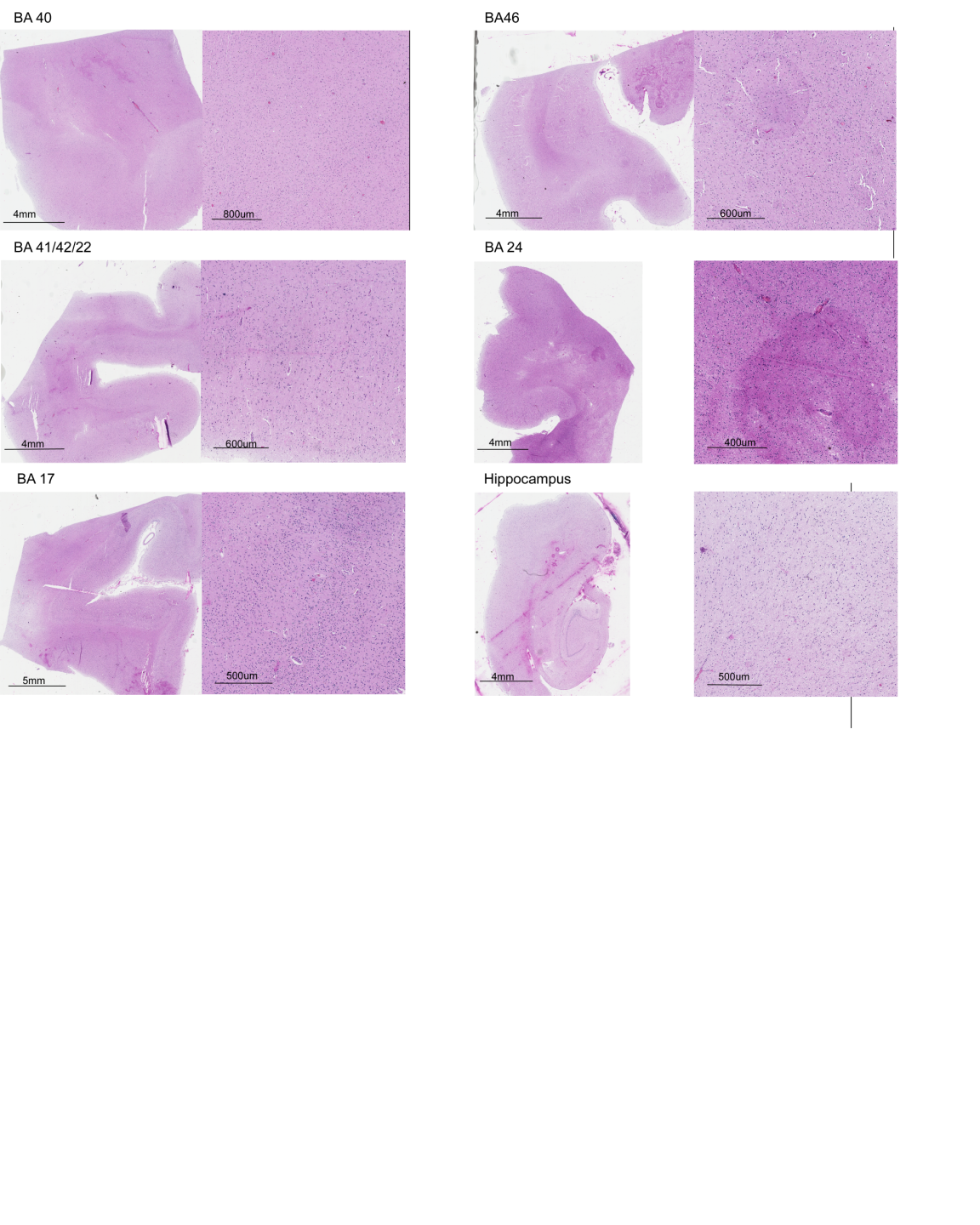

**Supplementary Figure 2: Reliability of iTRAQ analysis pipeline.** A: Technical CV distributions of label-free and iTRAQ quantification. B: The venn diagram of the proteins quantified in 4 batches. C: Log ratio distribution of iTRAQ quantification results. D: The ratio distributions of each run in 4 batches.

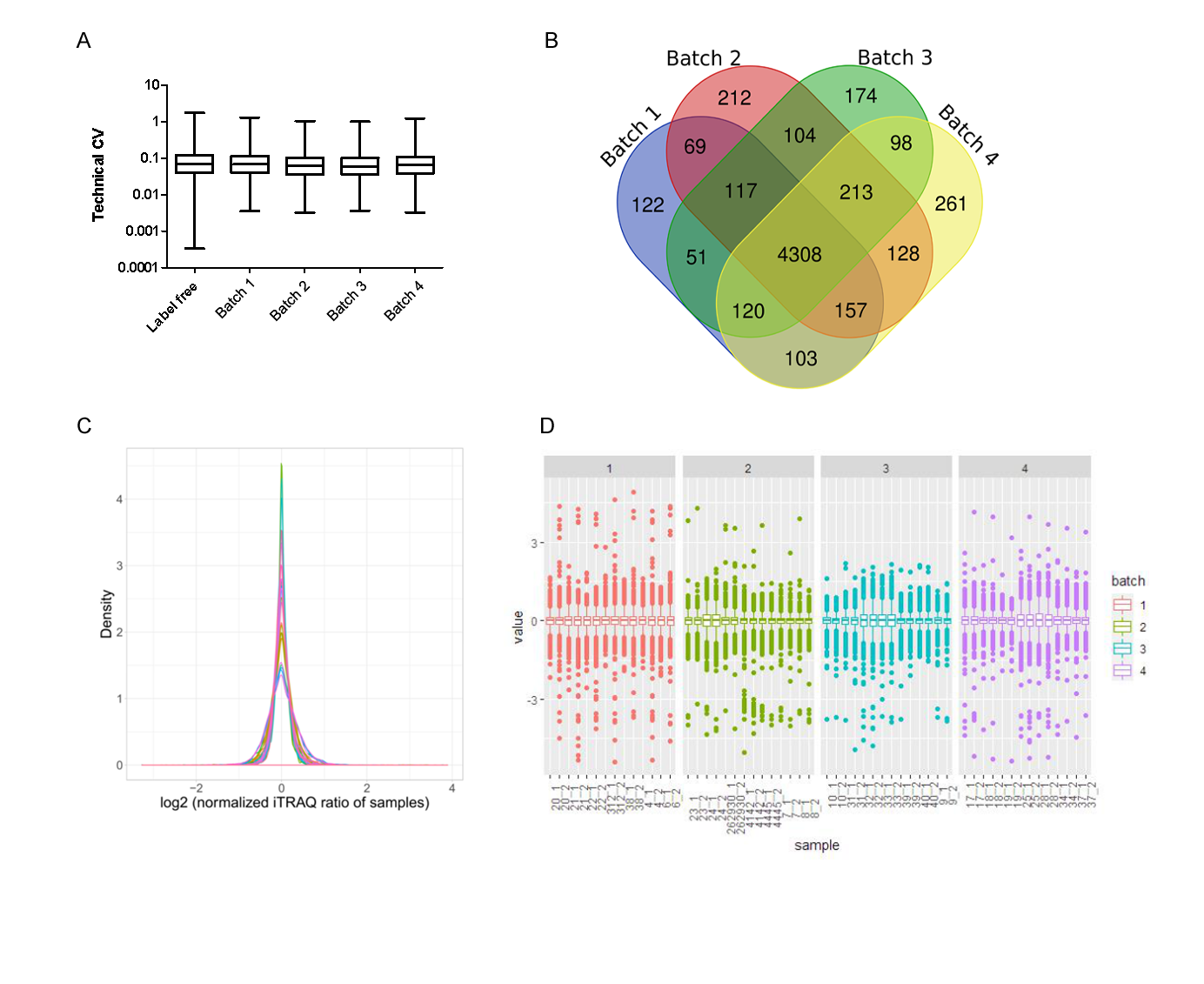

**Supplementary Figure 3: Protein abundance distribution of BA46 from the highest abundance to the lowest abundance protein.**

Log10 iBAQ

| Protein Rank | Log iBAQ |
| --- | --- |
| 1 | 9.48225 |
| 2 | 9.44469 |
| 3 | 9.34176 |
| 4 | 9.29338 |
| 5 | 9.26221 |
| 6 | 9.24366 |
| 7 | 9.19562 |
| 8 | 9.16573 |
| 9 | 9.13372 |
| 10 | 9.13292 |
| 11 | 9.05938 |
| 12 | 9.05111 |
| 13 | 8.98527 |
| 14 | 8.91639 |
| 15 | 8.7628 |
| 16 | 8.74595 |
| 17 | 8.72109 |
| 18 | 8.71118 |
| 19 | 8.70679 |
| 20 | 8.6694 |
| 21 | 8.64795 |
| 22 | 8.6375 |
| 23 | 8.60842 |
| 24 | 8.60405 |
| 25 | 8.60178 |
| 26 | 8.59472 |
| 27 | 8.58122 |
| 28 | 8.55795 |
| 29 | 8.55399 |
| 30 | 8.54754 |
| 31 | 8.54483 |
| 32 | 8.52986 |
| 33 | 8.51963 |
| 34 | 8.50661 |
| 35 | 8.50421 |
| 36 | 8.49683 |
| 37 | 8.4858 |
| 38 | 8.48318 |
| 39 | 8.47256 |
| 40 | 8.47188 |
| 41 | 8.45973 |
| 42 | 8.45869 |
| 43 | 8.45081 |
| 44 | 8.43351 |
| 45 | 8.43041 |
| 46 | 8.42725 |
| 47 | 8.42345 |
| 48 | 8.41183 |
| 49 | 8.40302 |
| 50 | 8.39993 |
| 51 | 8.39014 |
| 52 | 8.38248 |
| 53 | 8.37007 |
| 54 | 8.35165 |
| 55 | 8.34907 |
| 56 | 8.34352 |
| 57 | 8.33383 |
| 58 | 8.32175 |
| 59 | 8.31403 |
| 60 | 8.31095 |
| 61 | 8.30081 |
| 62 | 8.27986 |
| 63 | 8.27449 |
| 64 | 8.26616 |
| 65 | 8.26408 |
| 66 | 8.25827 |
| 67 | 8.24982 |
| 68 | 8.23797 |
| 69 | 8.2377 |
| 70 | 8.20418 |
| 71 | 8.19651 |
| 72 | 8.17763 |
| 73 | 8.17515 |
| 74 | 8.17256 |
| 75 | 8.17084 |
| 76 | 8.16894 |
| 77 | 8.16055 |
| 78 | 8.14756 |
| 79 | 8.1389 |
| 80 | 8.13115 |
| 81 | 8.12434 |
| 82 | 8.12103 |
| 83 | 8.11559 |
| 84 | 8.10468 |
| 85 | 8.101 |
| 86 | 8.09926 |
| 87 | 8.08909 |
| 88 | 8.08759 |
| 89 | 8.08151 |
| 90 | 8.07801 |
| 91 | 8.07371 |
| 92 | 8.06607 |
| 93 | 8.0651 |
| 94 | 8.06416 |
| 95 | 8.05796 |
| 96 | 8.05665 |
| 97 | 8.05554 |
| 98 | 8.05402 |
| 99 | 8.05242 |
| 100 | 8.04844 |
| 101 | 8.04285 |
| 102 | 8.04236 |
| 103 | 8.03774 |
| 104 | 8.03739 |
| 105 | 8.03314 |
| 106 | 8.03032 |
| 107 | 8.01259 |
| 108 | 8.00894 |
| 109 | 8.00768 |
| 110 | 8.00456 |
| 111 | 8.00241 |
| 112 | 8.00219 |
| 113 | 8.00028 |
| 114 | 7.9988 |
| 115 | 7.99382 |
| 116 | 7.99337 |
| 117 | 7.98489 |
| 118 | 7.97867 |
| 119 | 7.9753 |
| 120 | 7.9724 |
| 121 | 7.96817 |
| 122 | 7.96454 |
| 123 | 7.96438 |
| 124 | 7.96248 |
| 125 | 7.95211 |
| 126 | 7.92381 |
| 127 | 7.91762 |
| 128 | 7.90554 |
| 129 | 7.89321 |
| 130 | 7.8919 |
| 131 | 7.8804 |
| 132 | 7.87401 |
| 133 | 7.86048 |
| 134 | 7.86041 |
| 135 | 7.85629 |
| 136 | 7.85521 |
| 137 | 7.85398 |
| 138 | 7.85297 |
| 139 | 7.8489 |
| 140 | 7.8459 |
| 141 | 7.84399 |
| 142 | 7.84211 |
| 143 | 7.83851 |
| 144 | 7.83775 |
| 145 | 7.83449 |
| 146 | 7.83231 |
| 147 | 7.83148 |
| 148 | 7.82987 |
| 149 | 7.8296 |
| 150 | 7.82883 |
| 151 | 7.82819 |
| 152 | 7.82745 |
| 153 | 7.8272 |
| 154 | 7.82577 |
| 155 | 7.82564 |
| 156 | 7.82524 |
| 157 | 7.82306 |
| 158 | 7.82033 |
| 159 | 7.81915 |
| 160 | 7.81086 |
| 161 | 7.80832 |
| 162 | 7.80563 |
| 163 | 7.80297 |
| 164 | 7.80132 |
| 165 | 7.80064 |
| 166 | 7.8004 |
| 167 | 7.79954 |
| 168 | 7.79918 |
| 169 | 7.79608 |
| 170 | 7.79604 |
| 171 | 7.79514 |
| 172 | 7.79426 |
| 173 | 7.79304 |
| 174 | 7.7898 |
| 175 | 7.78871 |
| 176 | 7.78756 |
| 177 | 7.78328 |
| 178 | 7.78258 |
| 179 | 7.78113 |
| 180 | 7.78053 |
| 181 | 7.78002 |
| 182 | 7.77891 |
| 183 | 7.77837 |
| 184 | 7.77653 |
| 185 | 7.77459 |
| 186 | 7.77443 |
| 187 | 7.76064 |
| 188 | 7.76041 |
| 189 | 7.75874 |
| 190 | 7.75679 |
| 191 | 7.75556 |
| 192 | 7.74802 |
| 193 | 7.74784 |
| 194 | 7.74607 |
| 195 | 7.74388 |
| 196 | 7.74281 |
| 197 | 7.7425 |
| 198 | 7.74208 |
| 199 | 7.73795 |
| 200 | 7.73792 |

**Supplementary Figure 4: Data visualization in Human Brain Proteome Atlas website.** A: Protein quantification distribution of MAPT in 29 Brodmann areas. B: Protein expression data in Brodmann area 25 (Subgenual area). C: Comparison of the differential proteins between BA3/1/2, and BA4.

A

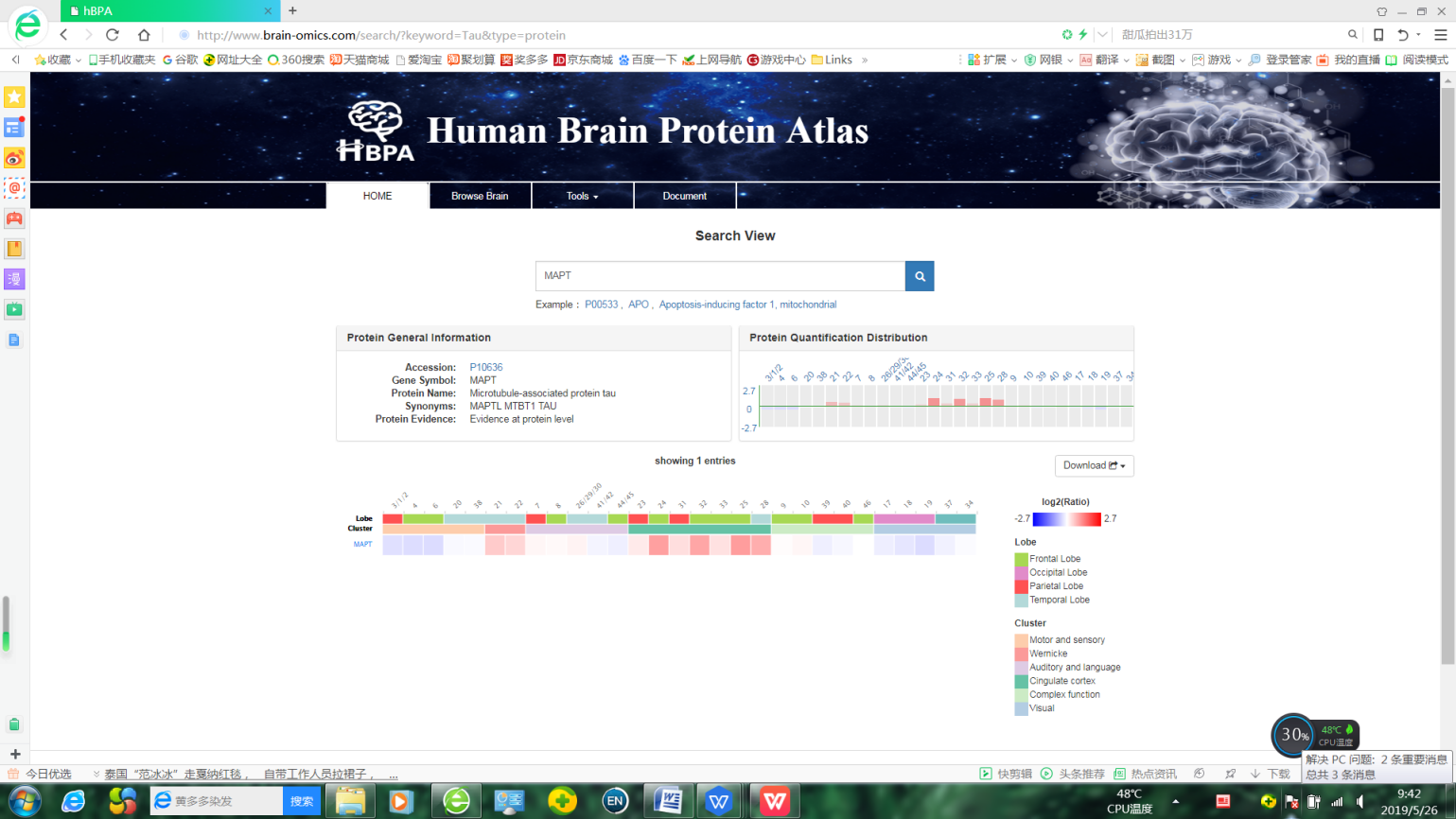

B

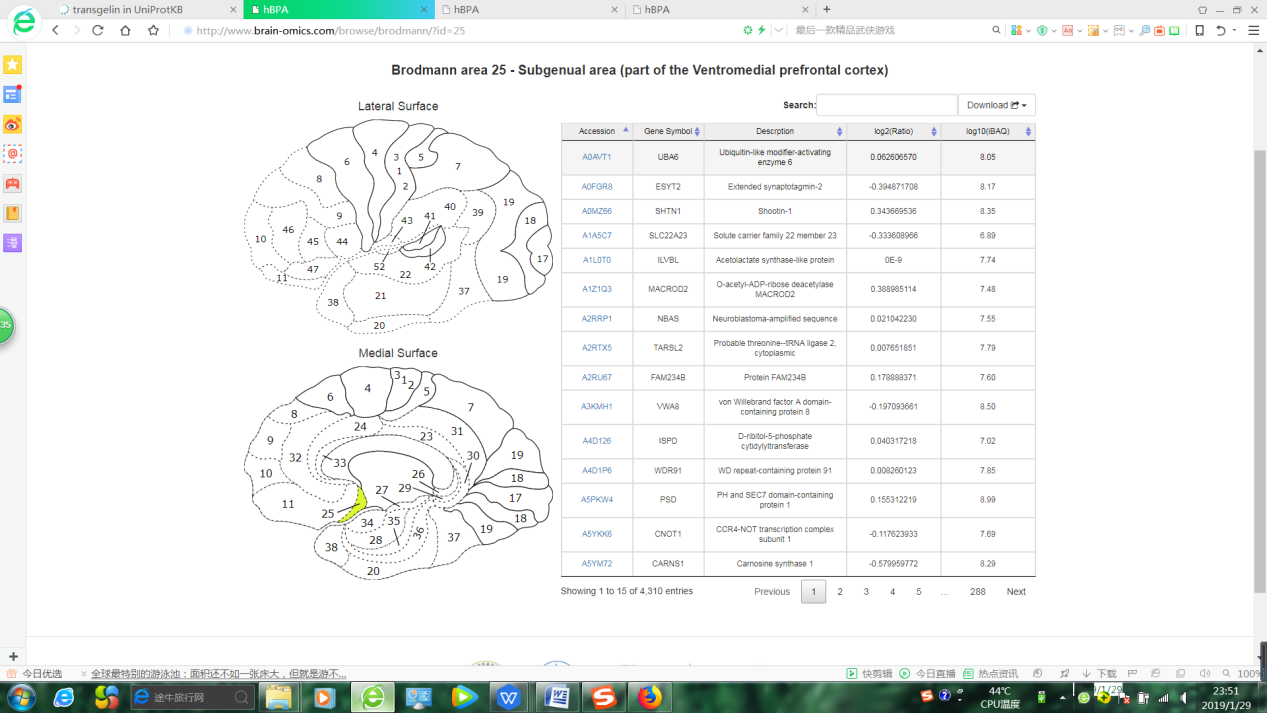

C

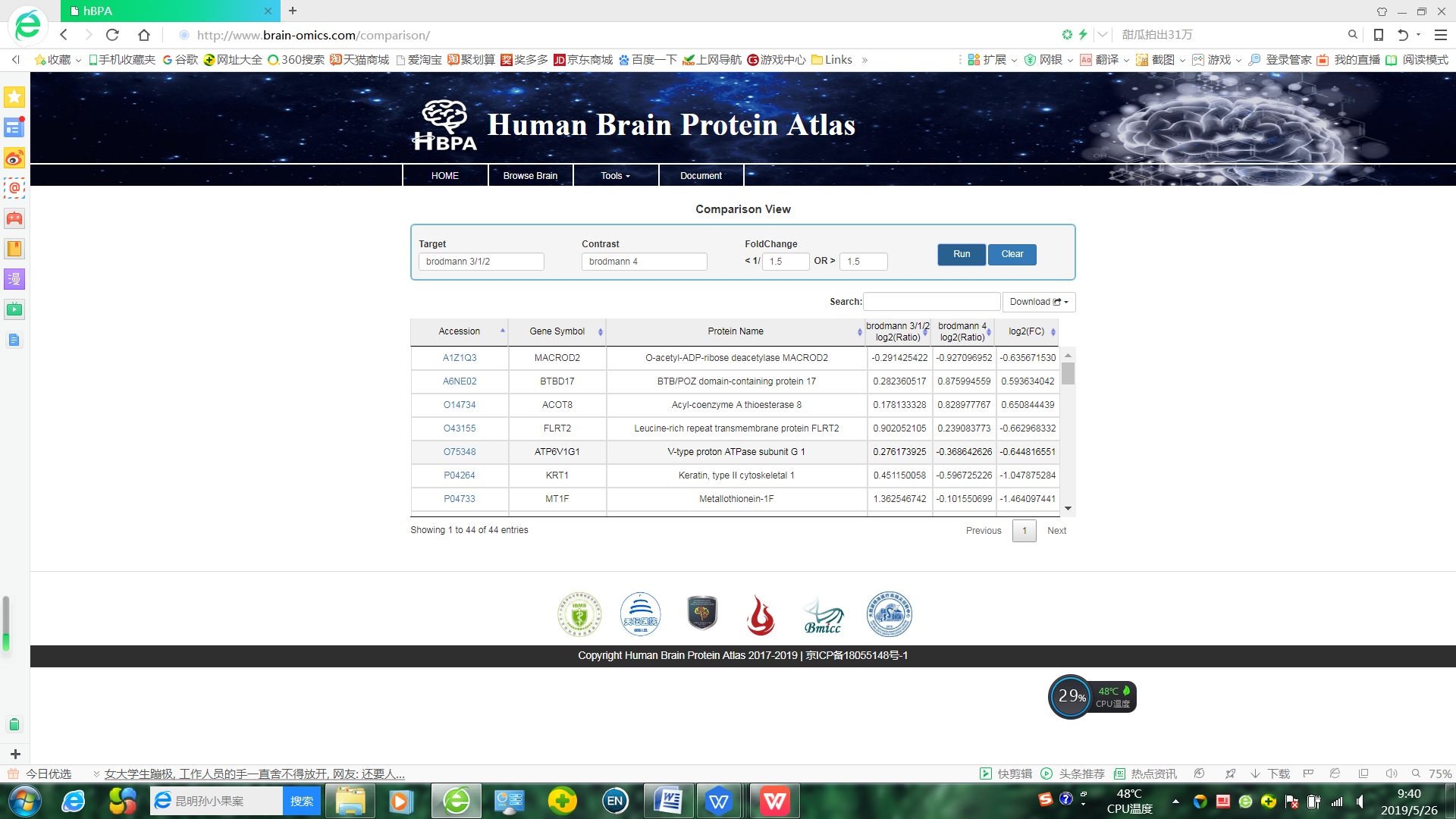

**Supplementary Figure 5: The protein clustering analysis.** A: The clustering is derived by consensus clustering based on 1,000 resampled data sets, exploring the range of from k=2 to k=8. B: Based on both consensus cumulative distribution function (CDF) area curve and the delta plot assessing change in CDF area, the CDF of consensus increased at greater k values. At k = 6, a relative increase in CDF reached a very small value. . C: Silhouette plots were performed to evaluate the Stability of the clustering. The silhouette plot at k = 6 showed a mean silhouette score of 0.72, and all scores were positive.

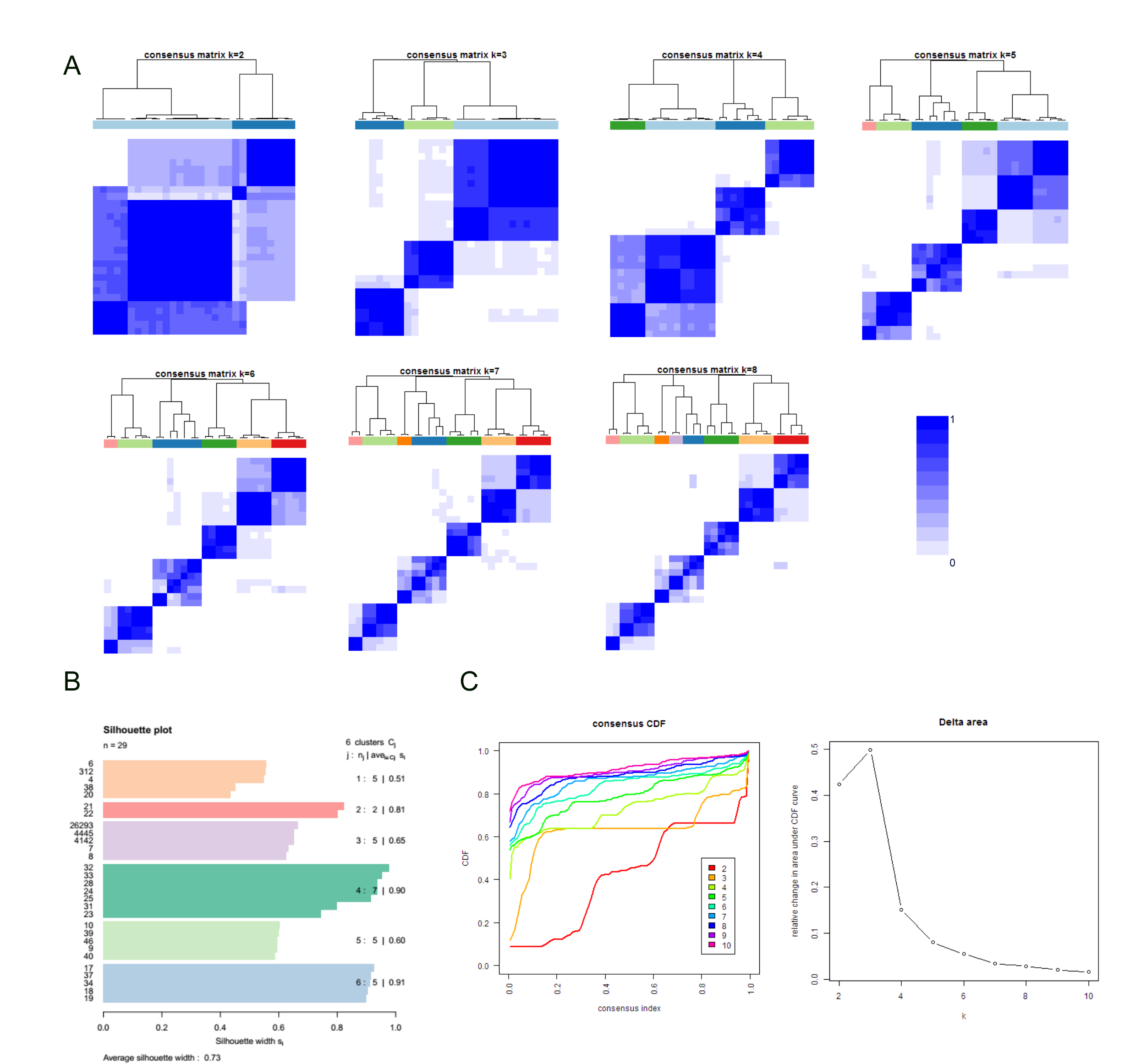

**Supplementary Figure 6: Analysis of the differential proteins between Wernicke’s area cluster and auditory and Broca’s area cluster.** A: IPA-based Canonical pathway analysis of differential proteins. The -Log (p-value) of the Canonical pathway term were shown. B: Heatmap of differential proteins involved in language eloquent regions related neurotransmitter pathways. The normalized protein expression value z-score represents the protein abundance. Red: high abundance. Blue: low abundance.

A

-Log(p-value)

|  | -log(p-value) |
| --- | --- |
| Actin Cytoskeleton Signaling | 3.21 |
| Sirtuin Signaling Pathway | 10.1 |
| Oxidative Phosphorylation | 16.3 |
| Mitochondrial Dysfunction | 17.8 |

B

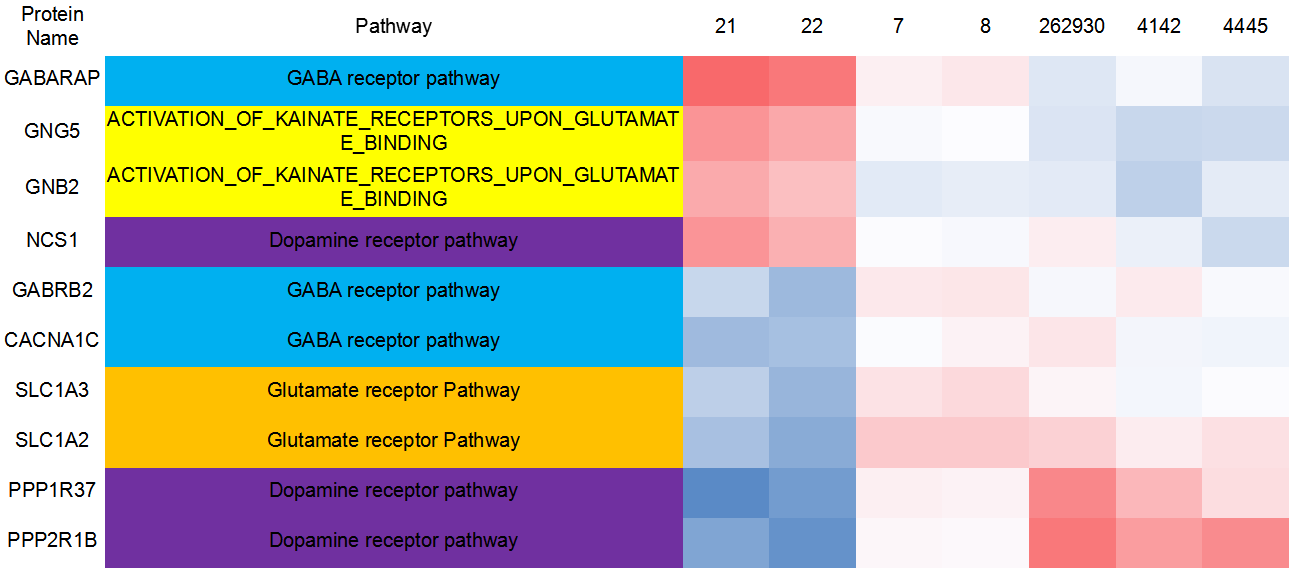

**Supplementary Figure 7: Functional analysis of cluster signature proteins, related to Figure 3.** A: The IPA-based biofunction (A) and KEGG database-based pathway (B) enrichment analysis of cluster signature proteins.The -Log (p-value) by two-sided hypergeometric test in heatmap. Red: significantly enriched. C-E: Heat map of the high-abundance proteins (C), mid-abundance proteins (D) and low abundance proteins (E) in the 29 BAs. The normalized protein expression value z-score represents the protein abundance. Red: high abundance. Blue: low abundance. F: Protein expression analysis of cingulate cortex cluster in Parkinson’s signaling. G: Protein expression analysis of cingulate cortex cluster in Alzheimer’s disease signaling. (Red: highly-expressed in cingulate cortex cluster; Green: lowly-expressed in cingulate cortex cluster)

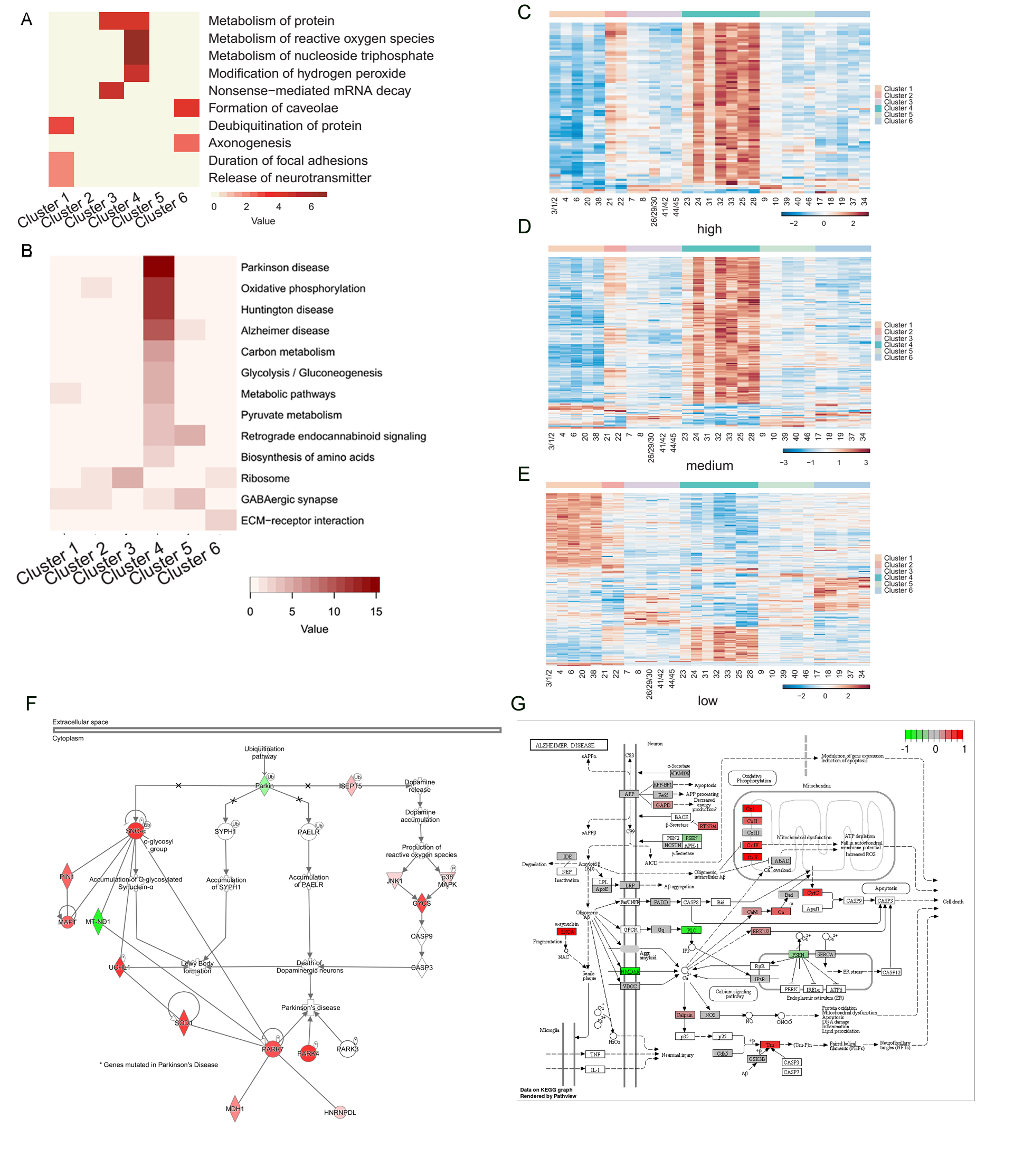

### Supplementary Figure 8: The Western Blot and IHC-Fr results of cluster signature proteins. A: Western Blot results of two motor and sensory cluster signature proteins (MRPS34 and EPM2A) and two cingulate cortex cluster signature proteins (Tau and a-syn). B: Comparison between immunoblotting results and iTRAQ results of above proteins. C: IHC-Fr validation results of two cingulate cortex cluster signature proteins (MAPT and SYUA). Scale bars represent as 50 µm.

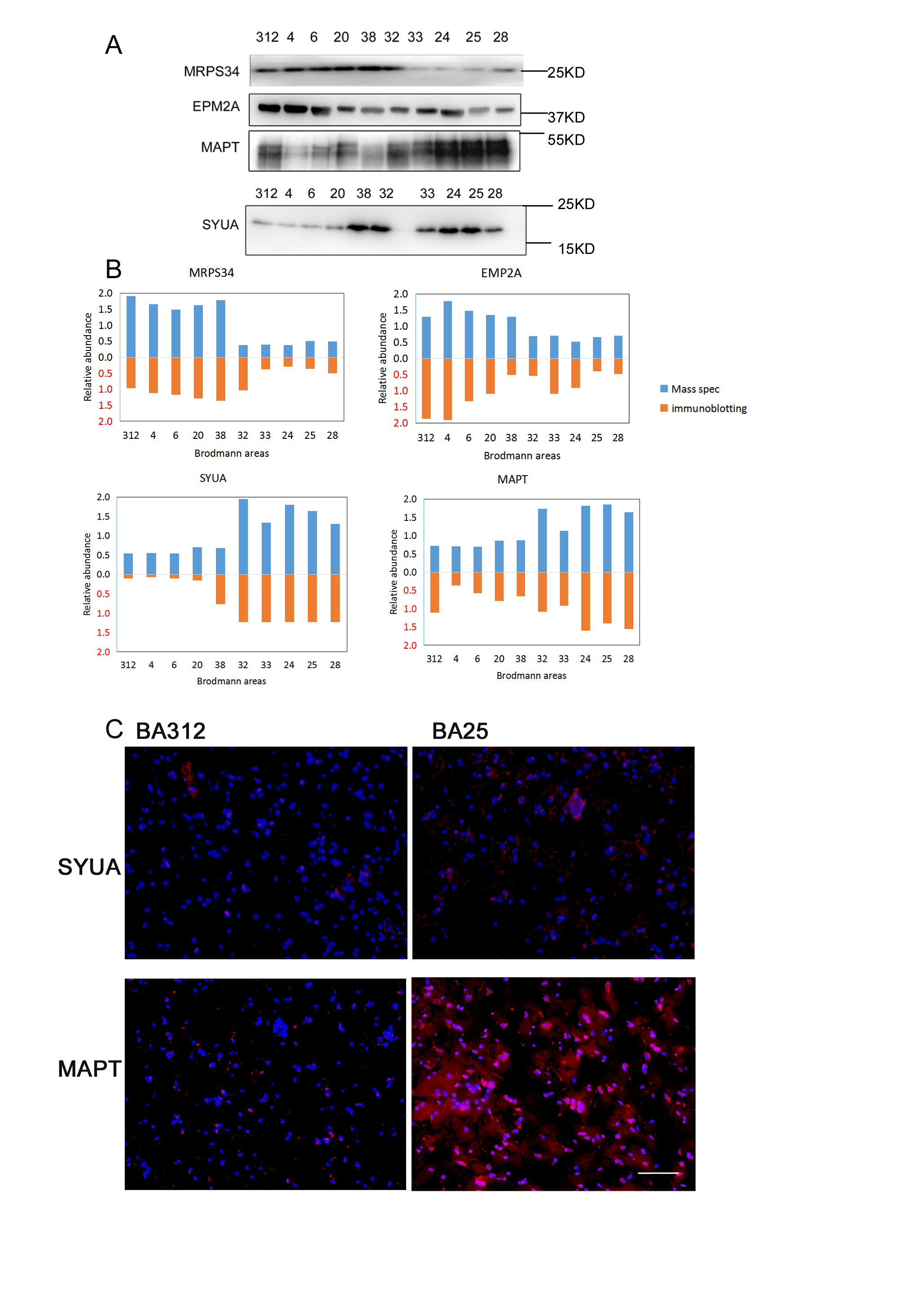

**Supplementary Figure 9: The Western Blot and IHC-Fr results of Brodmann area signature proteins.** A: Western Blot results of three Brodmann area signatureproteins (Transgelin, FAM107B, and RAD23A). B: Comparison between the immunoblotting results and iTRAQ results of above proteins.

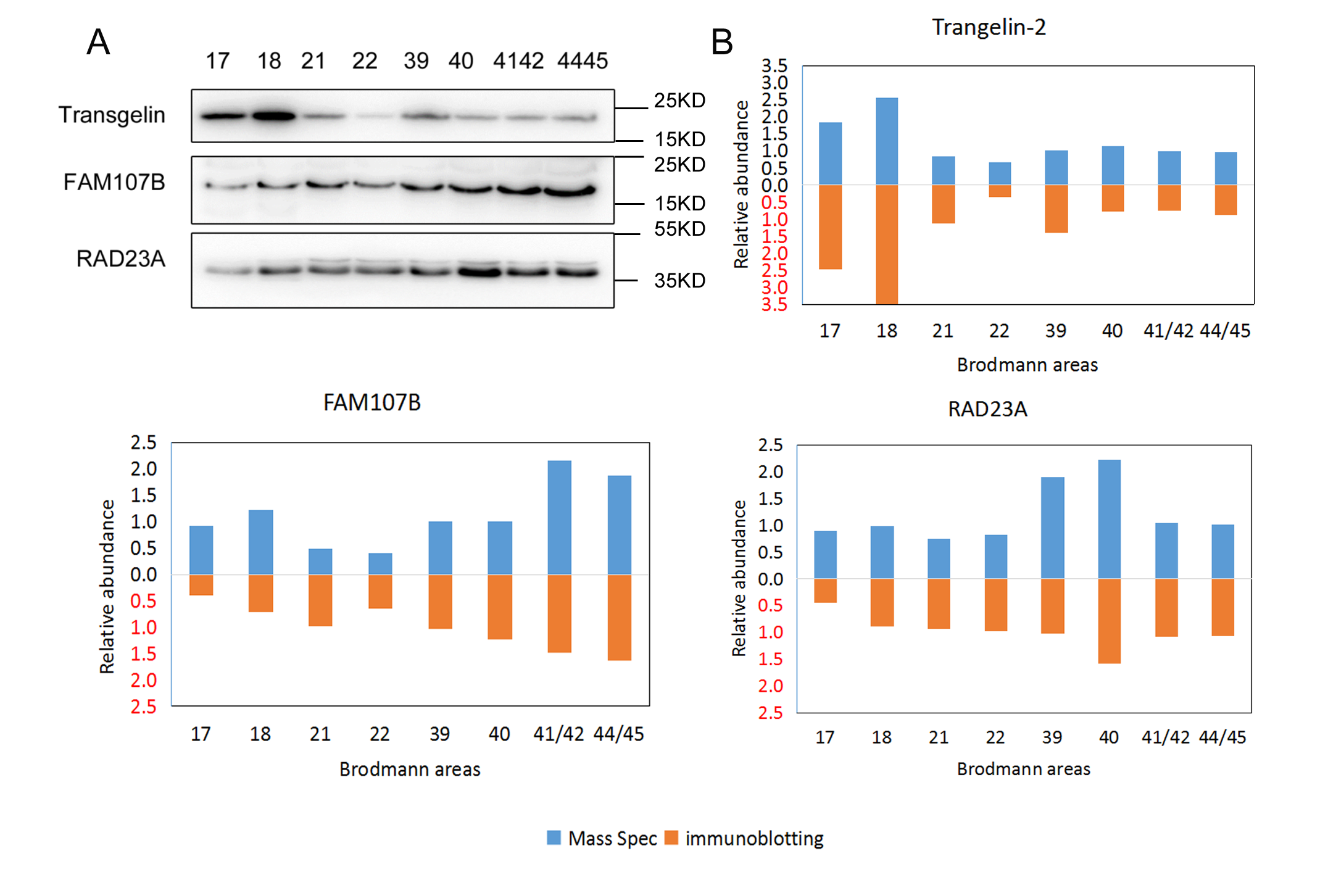

**Supplementary Figure 10: Quality control of RNA-seq.** A: Number of gene detected in RNA-seq analysis of the 21 BAs. B: Box plot of gene expression level of 21 BAs.

A

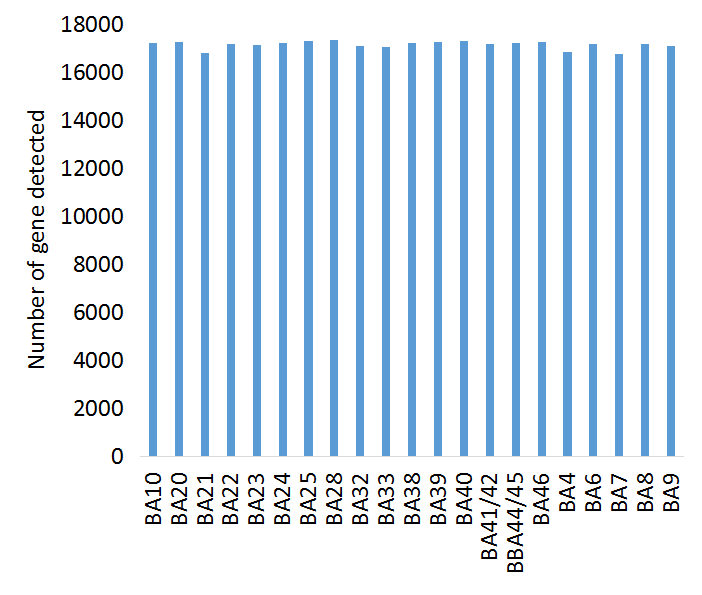

B

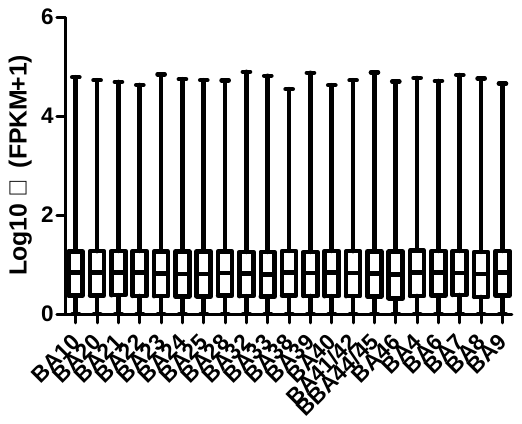

**Supplementary Figure 11: Scatter plots of mRNA and protein fold change of all regions compared with BA 46**. These scatter plots are identically defined as those in Figure 6E.

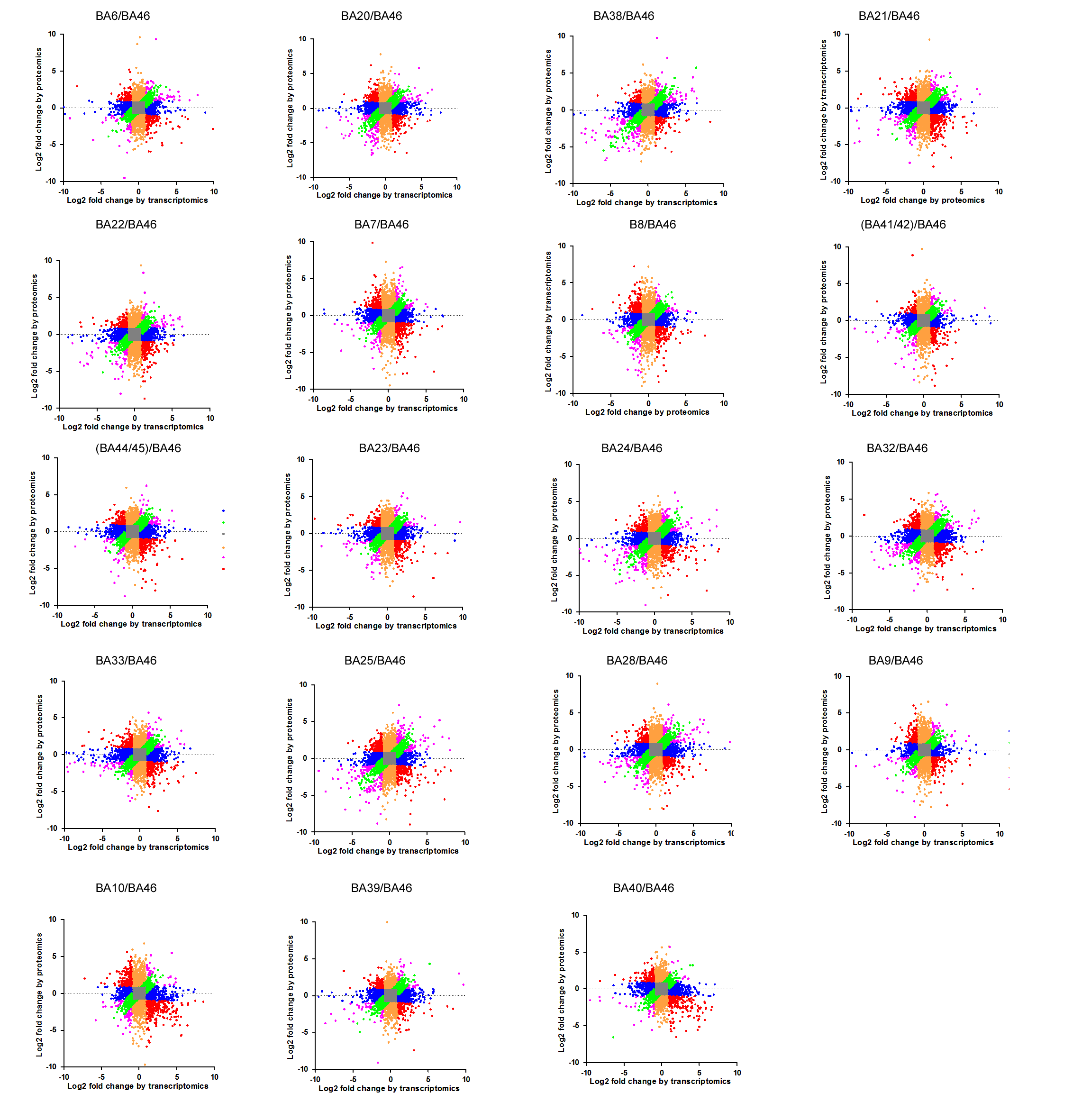
